# Glycan cross-feeding drives mutualism between *Fusobacterium* and the vaginal microbiota

**DOI:** 10.1101/463349

**Authors:** Kavita Agarwal, Lloyd Robinson, Justin Perry, Lynne Foster, Hueylie Lin, Brett Tortelli, Valerie P. O’Brien, Hilary Reno, Nicole Gilbert, Warren Lewis, Amanda Lewis

## Abstract

Dysbiosis of the vaginal microbiome is associated with vaginal colonization by potential pathogens including *Fusobacterium nucleatum*, a bacterium linked with intrauterine infection and preterm birth. However, mechanisms by which such pathogens gain a foothold in the dysbiotic vagina remain obscure. Here we demonstrate that sialidase activity, a biochemical marker of vaginal dysbiosis, promoted *F. nucleatum* foraging on mammalian sialoglycans, an otherwise inaccessible resource. In mice with sialidase-positive vaginal microbiomes, mutant *F. nucleatum* unable to consume sialic acids displayed impaired colonization. Furthermore, community- and co-culture experiments showed that *F. nucleatum* did not simply take advantage of sialidase-positive bacteria, but also “gave back” to the community, supporting robust outgrowth of sialidase-producers, including *Gardnerella vaginalis*. These results illustrate that mutualistic relationships between vaginal bacteria support pathogen colonization and reinforce dysbiosis, adding complexity to the simplistic dogma that the mere absence of “healthy” lactobacilli is what creates a permissive environment for pathogens during dysbiosis.

## Introduction

Bacterial vaginosis (BV) is a common vaginal dysbiosis occurring in approximately 1/3 of U.S. women (Allsworth and Peipert, 2007). It is characterized by low levels of “beneficial” lactobacilli and overgrowth of bacteria from diverse taxonomic groups (Hill et al., 1984). During pregnancy BV puts women at higher risk of preterm delivery and pregnancy complications caused by infection of the placenta and amniotic fluid (Goldenberg et al., 1996; Hillier et al., 1988; Hitti et al., 2001; Rezeberga et al., 2008; Silver et al., 1989). Although the condition is often asymptomatic, women with BV are more likely to acquire infections such as HIV, chlamydia, and gonorrhea than women without BV (Brotman et al., 2010; Peipert et al., 2008; Wiesenfeld et al., 2003). Additionally, BV puts women at higher risk of vaginal colonization by potential pathogens known to cause intrauterine infection. For example, BV-positive women are more likely to be vaginally colonized by *Fusobacterium nucleatum,* a Gram-negative spindle-shaped, obligate anaerobe, and one of the most commonly isolated microorganisms from amniotic fluid of women in preterm labor (DiGiulio, 2012; Han et al., 2009; Hill, 1998).

Unfortunately, mechanisms underlying the enhanced susceptibility to infection in BV are not well defined, due to the lack of tractable experimental mouse models, and limitations in culture-based approaches and genetic systems among fastidious vaginal anaerobes. A common feature of dysbiosis is the loss of “colonization resistance,” the ability of a stable healthy microbiome to thwart the entry of new potentially pathogenic members into the community. It is believed that a dominant mechanism behind the many infection-related health complications associated with BV is a permissive environment created by the lack of dominant lactobacilli in the vagina, which are known to produce antimicrobial molecules. We tested a different idea, based on the principle that ecosystems rely on mutually supporting metabolic interactions to stabilize communities. The bacterial communities of mucosal surfaces are no exception. Mechanistic examples applying this fundamental concept have so far been limited mainly to the gut (Belzer et al., 2017; Huang et al., 2015; Ng et al., 2013; Rakoff-Nahoum et al., 2016; Rey et al., 2013). Here we take advantage of models where dominant lactobacilli populations are absent, to test the hypothesis that establishment of potential pathogens such as *F. nucleatum* within the already occupied vaginal niche may depend on the ability to capitalize on nutrient resources uniquely available during vaginal dysbiosis.

One biochemical characteristic of the vagina during dysbiosis that is often used as a diagnostic feature of BV is the presence of sialidase activity in vaginal fluids (Briselden et al., 1992; McGregor et al., 1994; Myziuk et al., 2003). Sialidase activity may contribute to the complications of BV; for example high sialidase activity in vaginal fluids is associated with increased risk of preterm birth (Cauci and Culhane, 2011; McGregor et al., 1994). Sialidases play diverse mechanistic roles in bacterial-host interactions, co-infection, and dysbiosis in oral, gastrointestinal and airway systems (Huang et al., 2015; Kurniyati et al., 2013; Li et al., 2015; Lim et al., 2017; Mally et al., 2008; Ng et al., 2013; Siegel et al., 2014; Tailford et al., 2015; Wong et al., 2018). We hypothesize that bacterial sialidases also play important roles in pathogen colonization and dysbiosis in the vagina. Bacterial sialidases liberate abundant mucosal carbohydrate residues called sialic acids from glycan chains of secreted mucus components and cell surface glycoproteins, which can be used by some bacteria as a carbon source (Lewis and Lewis, 2012). Our earlier work showed that *Gardnerella vaginalis*, an abundant species in BV, secretes sialidase, which hydrolyzes sialic acids from host glycans in the extracellular space, followed by uptake and catabolism (Gilbert et al., 2013; Lewis et al., 2013). Consistent with these findings, vaginal mucus secretions of women with BV also have higher levels of free/liberated sialic acid (Lewis et al., 2013; Noecker et al., 2016; Srinivasan et al., 2015), and an overall depletion of total/bound sialic acids (Lewis et al., 2013; Moncla et al., 2015; Wang et al., 2015). Interestingly, not all bacteria that catabolize sialic acids express sialidases. Therefore, it is likely that the benefit of sialidases may extend beyond the sialidase producers to potential pathogens and/or other members of the dysbiotic vaginal microbiome.

Here, we asked whether the ability to metabolize sialic acids contributes to vaginal colonization by one potential pathogen, *F. nucleatum*, which we chose for three reasons. First, although better known for its roles in periodontal disease (Binder Gallimidi et al., 2015) and colorectal cancer (Zhang et al., 2018), *F. nucleatum* can also occupy the vagina in some women (Guerrero-Preston et al., 2017; Hitti et al., 2001; Holst et al., 1994). Second, *F. nucleatum* is more commonly found in the context of BV and less commonly in women with *Lactobacillus*- dominated vaginal microbiomes (Hitti et al., 2001; Holst et al., 1994). Third, *F. nucleatum* is one of the most common microorganisms identified in amniotic fluid of women in preterm labor (DiGiulio, 2012; Han et al., 2009), and its presence in vaginal fluids is a strong indicator of amniotic fluid infection among women in preterm labor (Hitti et al., 2001). Indeed, one study found that among pregnant women with BV, *Fusobacterium* was exclusively recovered from the vaginal samples of women who delivered preterm (Holst et al., 1994). So far, it is not known whether *F. nucleatum* benefits from or contributes to the growth of other organisms in the dysbiotic vaginal microbiome. We note that although some *F. nucleatum* strains seem to encode a sialic acid catabolic pathway (Caing-Carlsson et al., 2017; Gangi Setty et al., 2014; Manjunath et al., 2018; Yoneda et al., 2014), *F. nucleatum* is not known to express sialidases (Moncla et al., 1990). Thus, we wondered whether *F. nucleatum* might obtain sialic acids liberated by sialidases produced by other bacteria. Consistent with such an idea, *F. nucleatum* is found alongside sialidase-producing bacteria in the mouth and gut.

Here we investigate the cross-feeding of a specific metabolite, sialic acid, in the abundance and persistence of *F. nucleatum* in a mouse model of vaginal colonization. We show that *F. nucleatum* cannot on its own obtain sialic acid, but can benefit from sialic acid catabolism in the presence of sialidase-producing bacteria. We discovered that the capability to utilize sialic acids prolongs high titer *F. nucleatum* colonization in a vaginal niche containing a sialidase producing microbiota. In turn, *F. nucleatum* promotes the growth of sialidase-producing bacteria, both in mouse and human vaginal microbial communities. Together, our results may explain why women with BV are at increased risk of vaginal colonization by pathogens such as *F. nucleatum*. Additionally, our data suggest that mutual reinforcement between bacterial species, made possible through metabolite cross feeding, promotes pathogen colonization and vaginal dysbiosis.

## Results

### *F. nucleatum* catabolizes free sialic acid

To begin to determine whether *F. nucleatum* uses sialic acid catabolism to colonize the vagina, we first asked whether *F. nucleatum* encodes the necessary enzymes to utilize this carbon source. Some bacteria can import free sialic acids from the extracellular compartment via a sialic acid transporter (SiaT) and then use a sialate lyase (NanA) to convert them to *N*-acetyl-mannosamine (ManNAc) and phosphoenolpyruvate (Haines-Menges et al., 2015; Severi et al., 2007; Vimr and Troy, 1985). Recent studies suggest that at least some strains of *F. nucleatum* encode these functionalities (Gangi Setty et al., 2014; Manjunath et al., 2018; Yoneda et al., 2014). Our bioinformatic analysis of 28 *F. nucleatum* proteomes revealed that about half of the sequenced strains encoded a putative NanA (**Figure S1**). Homologs of NanA were identified in most members of the subspecies *nucleatum* and *vincentii*, [which includes the subspecies previously known as *fusiforme* (Kook et al., 2013)] and some strains of *animalis*. However, none of the *F. nucleatum* subsp. *polymorphum* strains encoded putative NanA homologs. Although NanA sequences in *F. nucleatum* strains showed low identity (~35%) to the sialic acid lyase from *Escherichia coli* (GenBank: AAC76257.1), they were highly conserved among the various strains and subspecies of *F. nucleatum.* In fact, they had greater than 95% identity to NanA from *F. nucleatum* subspecies *nucleatum* strain ATCC23726 (GenBank: EFG95907.1).

To determine whether the putative *F. nucleatum nanA* gene encodes a functional sialic acid lyase, we cloned it from ATCC23726 and generated a plasmid (pLR10; see **Table S1** for all plasmids and bacterial strains used in this study), which we transformed into a *nanA* deletion mutant of *E. coli* (strain MG1655) (Robinson et al., 2017). *E. coli* has a well-characterized sialic acid catabolic pathway (Vimr and Troy, 1985). *E. coli ΔnanA* was unable to grow on sialic acid as a sole carbon source when complemented with empty vector. However, the growth defect was rescued upon complementation with either *E. coli nanA* or *F. nucleatum nanA* (**Figure 1A**). In another set of experiments, we prepared lysates from these same *E. coli* strains and performed an assay for sialic acid lyase activity. Briefly, sialic acid (Neu5Ac) was incubated with clarified lysate supernatants, quenched at different time points, and Neu5Ac levels were quantified over time by DMB-HPLC as we have previously described (Lewis et al., 2007; Lewis et al., 2009; Lewis et al., 2006; Lewis et al., 2004). As expected, the *E. coli ΔnanA* strain containing empty vector did not have sialic acid lyase activity. However, in reactions with lysates from the complemented strains (*E. coli nanA* or *F. nucleatum nanA*) > 80% of the sialic acid was depleted within 30 minutes (**Figure 1B**). These results confirm that *F. nucleatum nanA* acts as a sialate lyase. Consistent with bioinformatic predictions, we confirmed that multiple strains of anaerobically cultivated *F. nucleatum* subspecies *nucleatum* consumed free sialic acid supplied in growth media, whereas the *polymorphum* and *animalis* strains tested did not (**Figure S1**). Analyses of other members of the *Fusobacteriales* suggest that some species in addition to *F. nucleatum* (e.g., *Fusobacterium mortiferum*) can catabolize sialic acids, but this is by no means a conserved feature of the *Fusobacteriales* (**Figure S1**).

**Figure 1.**
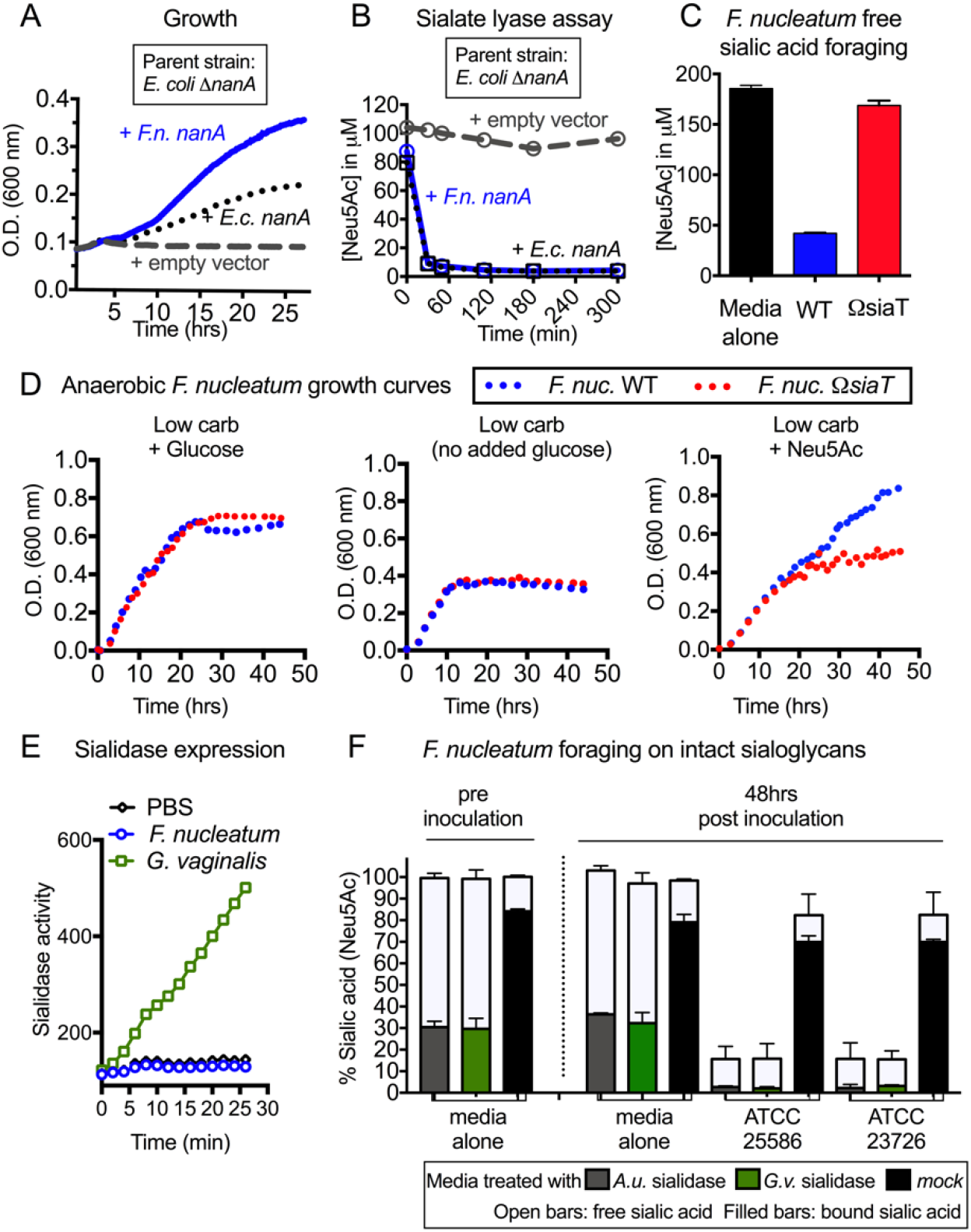
*F. nucleatum* can consume free sialic acid in the environment, but can only access sialic acids within glycans if sialidases from other bacteria are present. (A-B) *E. coli* MG1655 *ΔnanA* was complemented with empty vector, *E. coli nanA* or putative *nanA* from *F. nucleatum* 23726. *F.n.* = *F. nucleatum*; *E.c.* = *E. coli*. (A) Growth was assessed in minimal media with sialic acid by measuring absorbance at 600 nm. (B) Lysates of *E. coli nanA* mutant with empty vector or the complemented strain were incubated with Neu5Ac and its disappearance was monitored over time by DMB-HPLC. (C) Concentration of sialic acid (Neu5Ac) remaining in the medium 24 h post inoculation. Sialic acid consumption by *F. nucleatum* 23726 wild-type (WT) and *ΩsiaT* was studied in complete growth medium supplemented with free sialic acid. Data shown is representative of two independent experiments. Error bars show standard deviation from the mean value. (D) Growth of *F. nucleatum* (*F. nuc.)* WT and *ΩsiaT* in low carbohydrate (carb) medium with or without supplementation with the indicated carbohydrates. Data shown is representative of two independent experiments. (E) Cell associated sialidase activity of anaerobically cultured *G. vaginalis* JCP8151B and *F. nucleatum* ATCC23726 strains was analyzed using fluorogenic 4MU-Neu5Ac substrate. (F) *F. nucleatum* was grown anaerobically in media that was either untreated or exposed to purified sialidase from *Arthrobacter ureafaciens* (*A.u.)* or *G. vaginalis* (*G.v.).* Total and free sialic acid content was measured at 0 and 48 h post inoculation. Bound sialic acids (Bound = Total - Free) are inaccessible to *F. nucleatum*, except in the presence of exogenous sialidase. Data shown are representative of at least two technical replicates per condition performed in two or more independent experiments. Error bars represent standard deviation from the mean value. See also Figure S1, S2 and S3.

In *F. nucleatum*, the *nanA* gene is found downstream of a putative sialic acid transporter (*siaT*) in a gene cluster encoding other enzymes with predicted roles in carbohydrate metabolism (**Figure 2**, **Table S2**). To further assess the importance of *F. nucleatum* sialic acid catabolism, we constructed a minimal suicide vector targeting the *siaT* gene and used insertional mutagenesis to generate an *F. nucleatum* strain, *ΩsiaT,* in which *siaT* was disrupted (**Figure S2,** generated in the ATCC23726 SmR background). Hereafter, the *F. nucleatum* WT and *ΩsiaT* mutant are referred to as “WT” and “*ΩsiaT”.* When *ΩsiaT* was inoculated into growth media containing free sialic acid (Neu5Ac), the bacteria were unable to consume the free sialic acid, unlike the WT strain (**Figure 1C**). To further investigate the impact of sialic acid foraging on the growth of *F. nucleatum*, we inoculated WT and *ΩsiaT* into NYCIII media that was low in carbohydrates (no added glucose), contained glucose, or contained free sialic acid (Neu5Ac). WT and *ΩsiaT* grew similarly in media containing glucose, both plateauing at an OD_600_ of ~0.6. In low-carbohydrate media, both strains grew less well, plateauing at an OD_600_ of ~0.4. In contrast, in media containing sialic acid as the main source of free carbohydrate, *ΩsiaT* showed a substantial growth defect, reaching an OD_600_ of only ~0.4, whereas WT grew to an OD_600_ of ~0.8 (**Figure 1D**). These data confirm the genetic basis for sialic acid catabolism in *F. nucleatum* and show that it can benefit from its ability to catabolize free sialic acid under low glucose conditions.

**Figure 2.**
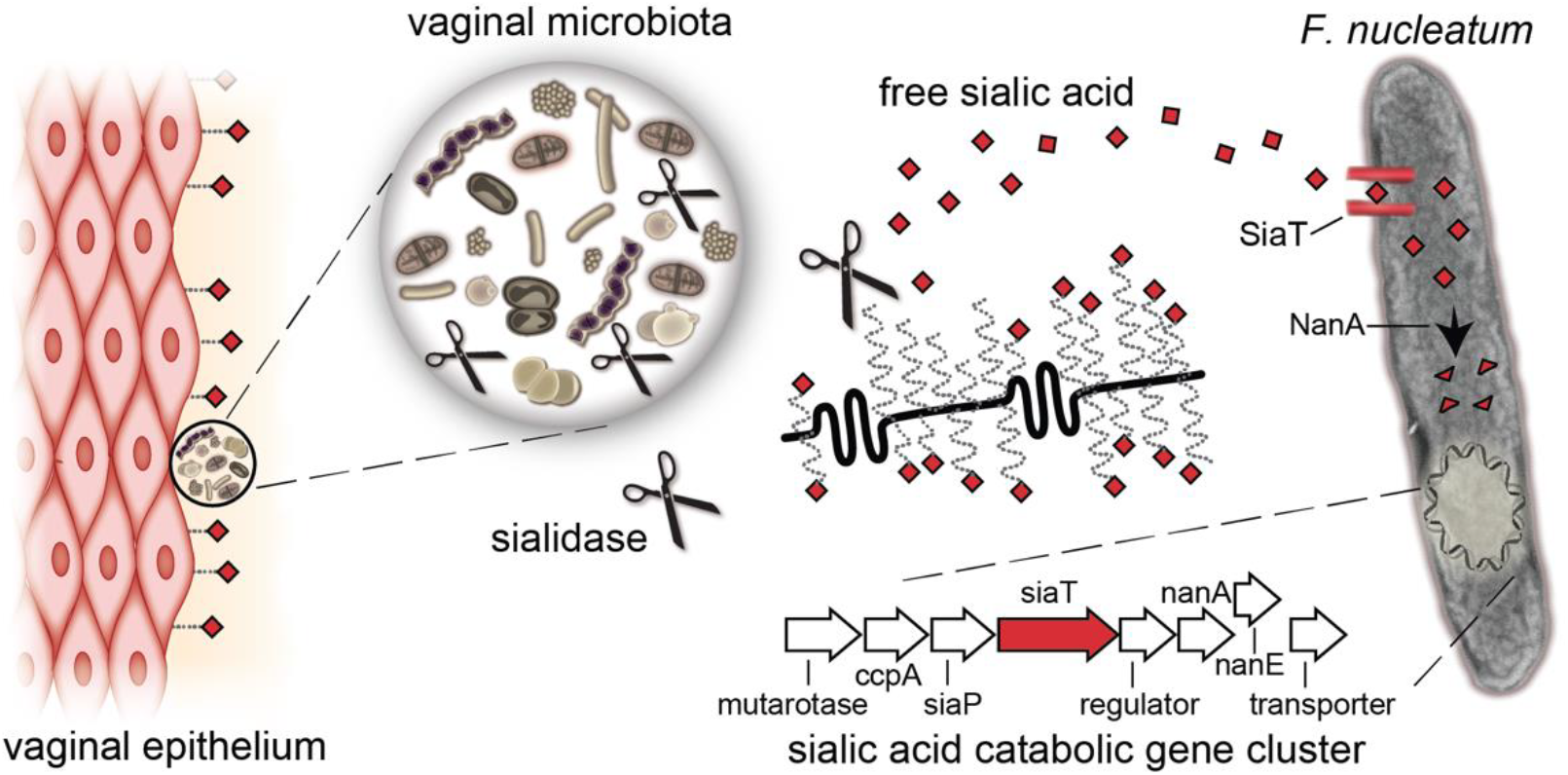
*F. nucleatum* 23726 takes up and catabolizes free sialic acid released by exogenous sialidases. Sialidase-producers in the vaginal microbial community release free sialic acids (red diamonds) from host glyco-conjugates, which may be accessed by *F. nucleatum*, which does not produce sialidase. SiaT = sialic acid transporter; NanA = *N*-acetylneuraminate lyase; More information about *F. nucleatum* genes shown in sialic acid catabolic gene cluster, and the enzymes they encode, can be found in Table S2.

### *F. nucleatum* accesses bound sialic acid from glycan chains, but only when liberated by exogenous sialidases

We next set out to determine whether any *F. nucleatum* strains encoded a potential sialidase, which is necessary to liberate sialic acids from glycan chains. Our bioinformatic analysis of *F. nucleatum* proteomes revealed no putative sialidases in this taxon. This is consistent with a previous study showing that 39 cultured *F. nucleatum* isolates all lacked sialidase activity (Moncla et al., 1990). In an *in vitro* assay, using the known sialidase producer *G. vaginalis* as a positive control, we show that anaerobically grown *F. nucleatum* could not cleave a fluorogenic sialic acid substrate (**Figure 1E**). Along with previous findings, these results establish that *F. nucleatum* lacks sialidase activity.

Given that *F. nucleatum* lacks its own sialidase, we hypothesized that its ability to consume glycan-bound sialic acids would require exogenous sialidase. To test this, growth media was treated overnight (O/N) with commercial recombinant purified sialidase from *Arthrobacter ureafaciens* (*A.u.),* partially purified *G. vaginalis* sialidase, or buffer alone. As expected, media treated with sialidases contained more free sialic acid than mock-treated culture media, in which sialic acids remained mostly in the bound form (**Figure 1F**). We then inoculated each media condition with *F. nucleatum*. After 48 hours, total sialic acids were significantly depleted only in media containing exogenous sialidase (**Figure 1F**). Similar results were obtained in experiments using sialidase from *Prevotella bivia* (strain ATCC29303), another BV-associated bacterium (**Figure S3**). Together, these data demonstrate that *F. nucleatum* can access, import, and consume sialic acids from sialo-glycoconjugates, but only if sialidase activity from an exogenous source is available to first liberate sialic acids into the free form (see schematic in **Figure 2**).

### C57BL/6 mice from different vendors have distinct vaginal microbiomes and variation in endogenous vaginal sialidase activity

Given our *in vitro* findings, and the fact that women with BV are unique in having sialidase-producing vaginal microbiomes, we wanted to determine whether sialic acid catabolism facilitates *F. nucleatum* colonization in a sialidase-positive vaginal environment. To answer this question, we first sought to identify a mouse model that harbored a sialidase-producing vaginal microbiota. We thus measured sialidase activity in vaginal washes from C57BL/6 mice purchased from multiple vendors. Endogenous vaginal sialidase activity was present in most mice from Envigo but was rarely detectable in mice from three other vendors (**Figure 3A**). Furthermore, Envigo mice had even more vaginal sialidase activity after treatment with β-estradiol, whereas mice from Jackson were sialidase-negative with or without β-estradiol treatment (**Figure 3B**). This is significant because our mouse model, like many other bacterial vaginal inoculation models (de Jonge et al., 2011; Gilbert et al., 2013; Jerse, 1999; Patras and Doran, 2016), employs β-estradiol treatment to maintain a state of pseudo-estrus. Estrogenized mice from Envigo were consistently sialidase-positive in four independent studies of mice purchased over two years (data not shown).

**Figure 3.**
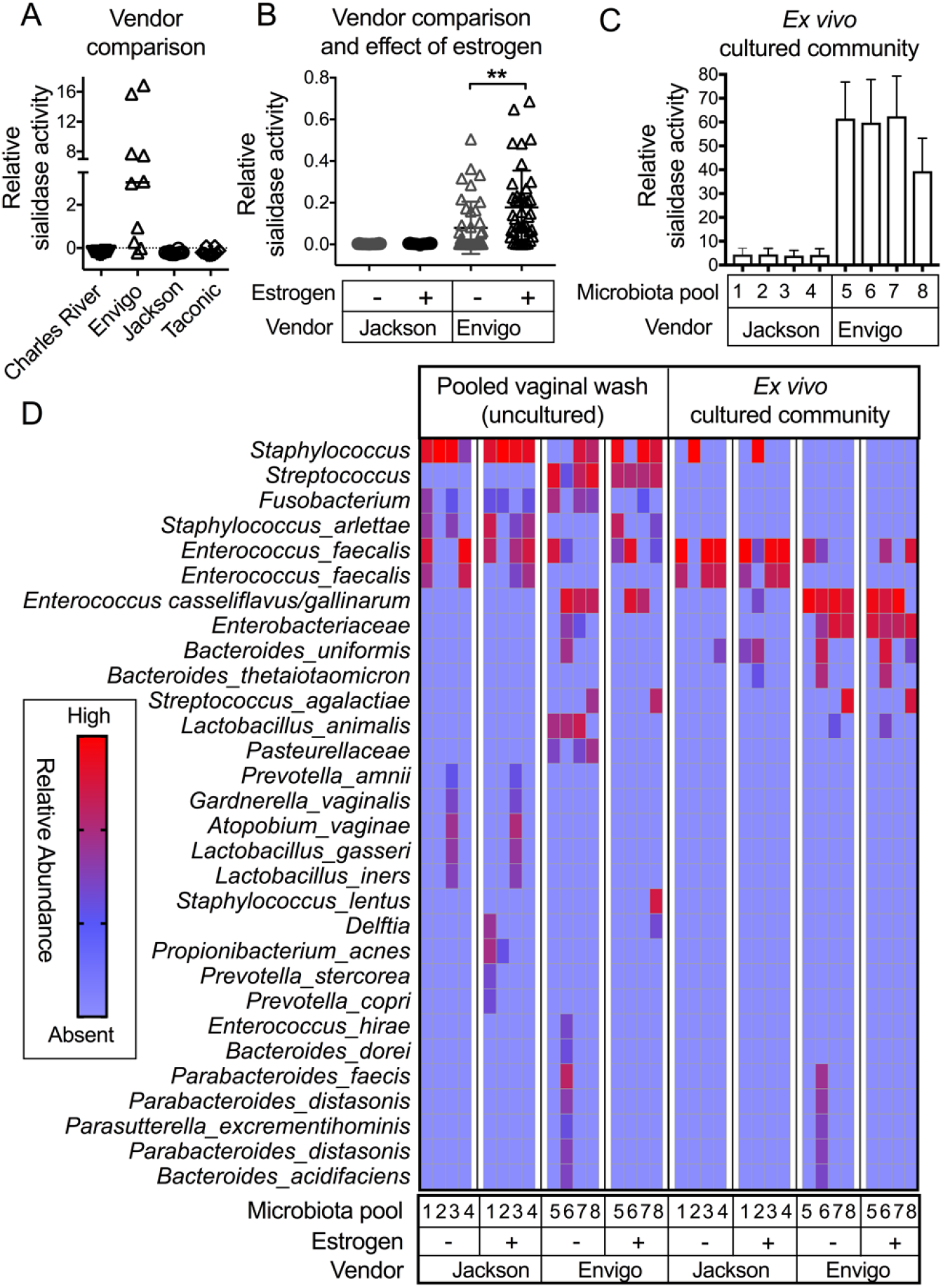
C57BL/6 mice from Envigo have endogenous vaginal sialidase activity. (A) Sialidase activity in vaginal washes of mice (not estrogenized) from different vendors measured using the fluorogenic 4MU-Neu5Ac substrate. N = 10 mice/ vendor. (B) Vaginal sialidase activity is elevated at 72 h post-estrogenization in mice from Envigo. No sialidase activity was detected in vaginal washes of mice from Jackson. N = 40 mice/ vendor. Wilcoxon paired-sign rank test was used for pairwise comparison of sialidase activity before and after estrogen treatment. ** *P* < 0.01. (C) Sialidase activity in microbiota pools from Envigo and Jackson mice, collected after estrogenization. Each “microbiota pool” consists of a cultured vaginal community from pooled vaginal wash of 4-5 co-housed mice. For A, B and C - Data shown is combined from two independent biological replicates. (D) Heat map shows abundance of different OTUs in the vaginal wash specimens of Envigo and Jackson mice collected before and after estrogenization. Microbiome analysis was done on uncultured and cultured vaginal washes pooled from 5 mice housed in the same cage. Each column represents one pool = 1 cage = 5 mice, total = 4 specimens per pooled condition. OTU = Operational taxonomic units. OTUs were assigned using UPARSE-OTU algorithm. Data was clustered using a hierarchical Euclidean method. See also Figure S4.

The finding that Envigo mice had sialidase activity in their vaginas, but Jackson mice did not, led us to wonder whether the vaginal microbiomes differed in the two groups of mice. To investigate whether sialidase activity in vaginal washes was of bacterial origin, and to generate sufficient material for such studies, we pooled vaginal washes from all five mice in each of four cages from each vendor. A portion of each vaginal wash pool was cultured anaerobically and frozen in cryoprotectant. We then extracted DNA and performed 16S community profiling on both 1) the original (<u>uncultured)</u> pooled wash material, and 2) the viable communities of bacteria (microbiota pools) derived from the <u>cultured</u> vaginal wash material (**Schematic S1**). Corroborating our findings in individual vaginal washes, microbiota pools from Envigo mice, but not Jackson mice, had high sialidase activity (**Figure 3C**). Microbiome profiles from samples before and after estrogen treatment had similar patterns of taxa, regardless of vendor. Since not all members of the microbiome will be cultivated under a single condition, the cultured and uncultured vaginal material had different proportions of some taxa (**Figure 3D and Figure S4**). Consistent with differences in sialidase activity, community profiling revealed that Envigo mice (sialidase-positive) had different vaginal microbiotas than Jackson mice (sialidase-negative). For example, several of the uncultured vaginal wash pools and *ex vivo* cultured communities from Envigo mice contained *Enterococcus casseliflavus/gallinarum*, whereas the corresponding samples from Jackson mice did not (**Figure 3D and Figure S4**). Colonies of *E. gallinarum* and *Bacteroides spp*. from Envigo mice expressed sialidase activity (data not shown). Together, these findings establish that C57BL/6 mice from Envigo exhibit vaginal sialidase activity arising from the endogenous microbiome.

### Sialic acid catabolism prolongs *F. nucleatum* vaginal colonization in animals with a sialidase-positive vaginal microbiome

We next wanted to determine whether *F. nucleatum* could use its ability to catabolize sialic acid to colonize the sialidase-positive Envigo vaginal niche. Thus, mice from Envigo and Jackson were vaginally inoculated with *F. nucleatum* WT or Ω*siaT*, and vaginal washes were collected longitudinally for several weeks. Microbiome profiling of pooled washes (from all five inhabitants of each cage) at 1-day post-inoculation (dpi) revealed that *F. nucleatum* became one of the dominant members of the vaginal microbiota (**Figure 4A**) in mice from both the vendors. We next compared the ability of WT and *ΩsiaT F. nucleatum* to colonize Envigo mice and found that *ΩsiaT* was at significantly lower titers than WT as early as 72 hours post inoculation (**Figure 4B and Figure S5**). However, at this time point, there was no difference in colonization between WT and *ΩsiaT* in sialidase-negative Jackson mice (**Figure 4C and Figure S5**).

**Figure 4.**
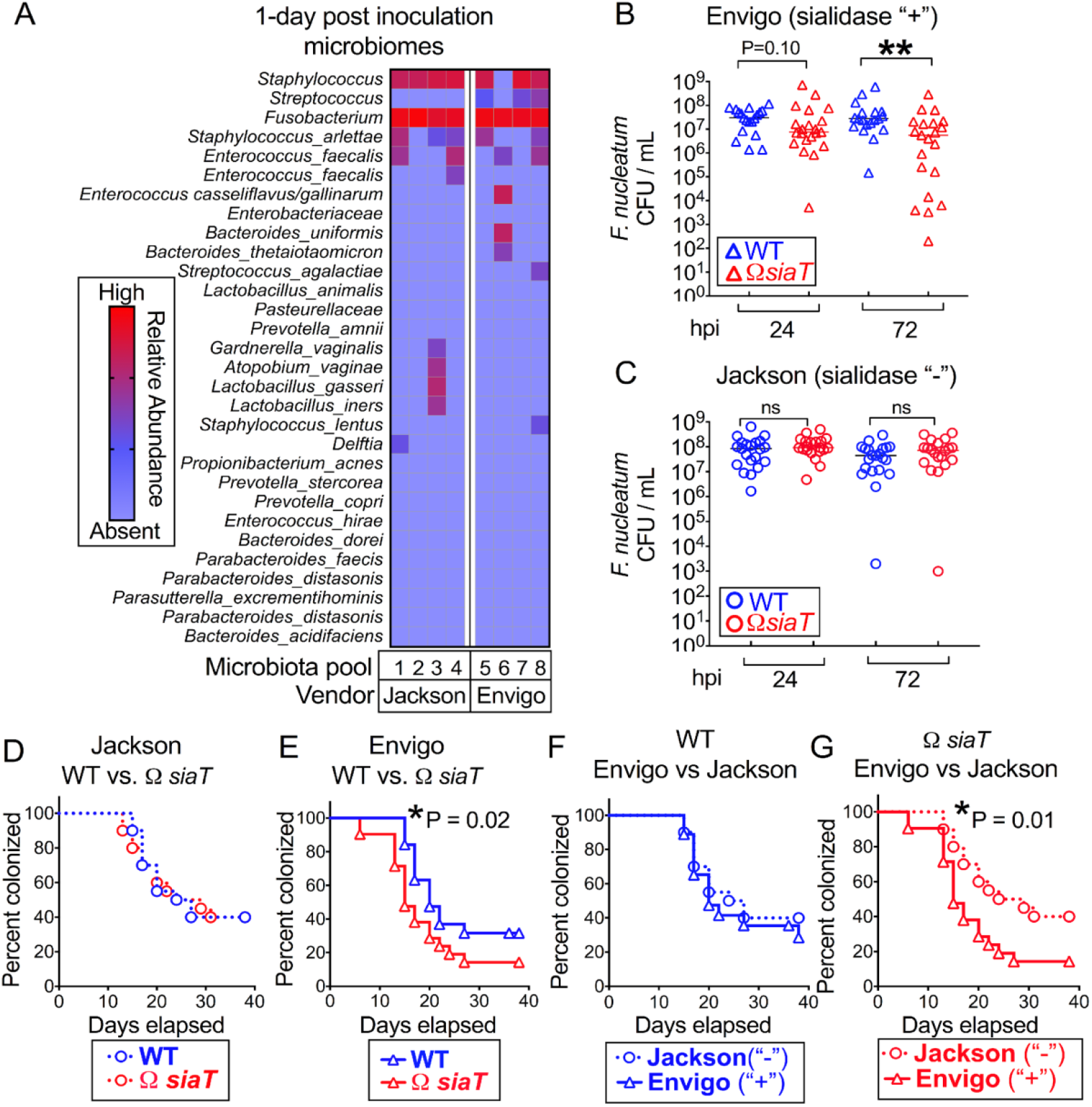
Sialic acid catabolism facilitates persistent *F. nucleatum* vaginal colonization in the setting of an endogenous sialidase-producing microbiome. (A) Heat map shows abundance of OTUs in uncultured pooled vaginal specimens (1 pool = 1 cage = 5 mice) collected 1-day post inoculation with *F. nucleatum*. OTUs were assigned using the UPARSE-OTU algorithm. Data was clustered using hierarchical Euclidean clustering. (B, C) *F. nucleatum* titers in vaginal wash collected at indicated time points post inoculation (B) in mice from Envigo, and (C) in mice from Jackson. ** *P* < 0.01, Mann-Whitney. (D-G) Course of C57BL/6 mice (Envigo and Jackson) colonization with the *F. nucleatum* wild-type (WT) and *ΩsiaT* mutant. Number of mice colonized in percent (*y* axis) was monitored on day 1 and every 2 days for 38 days (x axis). Statistically significance assessed by Log Rank test, * *P* < 0.05. (D) Comparison of WT vs. *ΩsiaT* colonization in Jackson mice. (E) Comparison of WT vs. *ΩsiaT* colonization in Envigo mice. (F) Comparison of WT colonization in Envigo vs. Jackson mice. (G) Comparison of *ΩsiaT* colonization in Envigo vs. Jackson mice. For Kaplan-Meier analysis, mice were considered cleared when no cfu were detected in undiluted wash at 2 consecutive time points. The graphs represent combined data from two experiments performed separately. OTU = operational taxonomic unit, hpi = hours post inoculation. See also Figure S5.

Following *F. nucleatum* colonization over several weeks, we found that, WT and Ω*siaT* remained at high levels in sialidase-negative Jackson mice through ~ 15 dpi, then gradually diminished at similar rates, both reaching 50% of mice still carrying WT or Ω*siaT F. nucleatum* at ~22 dpi (**Figure 4D**). In contrast, when inoculated into sialidase-positive Envigo mice, Ω*siaT* clears earlier than WT and is eliminated from a greater proportion of animals at any given time point over the duration of the experiment (P = 0.02, **Figure 4E**). The prolonged colonization of WT *F. nucleatum* in Envigo mice suggests that the ability to take up and catabolize sialic acid sustains *F. nucleatum* vaginal colonization in the sialidase-positive environment. Comparing WT and *ΩsiaT* colonization in different types of vaginal environments, we observed no significant difference in persistence of WT *F. nucleatum* between Envigo and Jackson mice (**Figure 4F**). However, in the sialidase-positive environment of Envigo mice, *ΩsiaT* was cleared significantly faster in contrast to persistent long-term colonization observed in Jackson mice (similar to WT *F. nucleatum*) (**Figure 4G**). We thus conclude that the ability to take up and catabolize sialic acid promotes *F. nucleatum* vaginal colonization in a sialidase-positive environment but is dispensable in a sialidase-negative environment.

### Sialidase activity from the mouse microbiome is sustained by colonization of *F. nucleatum*

While working with the murine system, we observed that, once received in our facility, vaginal sialidase activity waned rapidly in mice not inoculated with *F. nucleatum* (data not shown), or inoculated with a different urogenital pathogen such as *E. coli* (**Figure S6**). In contrast, when inoculated with *F. nucleatum*, vaginal sialidase activity was maintained at higher levels for a longer duration (**Figure S6**). This data suggested that *F. nucleatum* may be supporting the endogenous sialidase-producing vaginal community. Having discovered that *F. nucleatum* benefited from its ability to catabolize sialic acids in the presence of sialidase producing vaginal microbiota, we wondered whether the opposite was true. Does *F. nucleatum* also support the endogenous sialidase-producing vaginal community? And if so, does the capacity to catabolize sialic acids influence reciprocation by *F. nucleatum*? To test this idea, we first compared sialidase activity over time in mice inoculated with WT or *ΩsiaT F. nucleatum.* Interestingly, inoculation of WT *F. nucleatum* helped to maintain higher levels of vaginal sialidase for a longer duration as compared to inoculation with *ΩsiaT* (**Figure 5A**). While sialidase activity was equivalent between WT and *ΩsiaT*-inoculated mice at 1 dpi (**Figure 5B**), by 6 dpi sialidase activity was significantly lower in vaginal washes from mice inoculated with *ΩsiaT* compared to WT. These data suggest that colonization by WT sialic-acid consuming *F. nucleatum* helps sustain the production of sialidase by other bacteria in the vaginal microbiota of Envigo mice.

**Figure 5.**
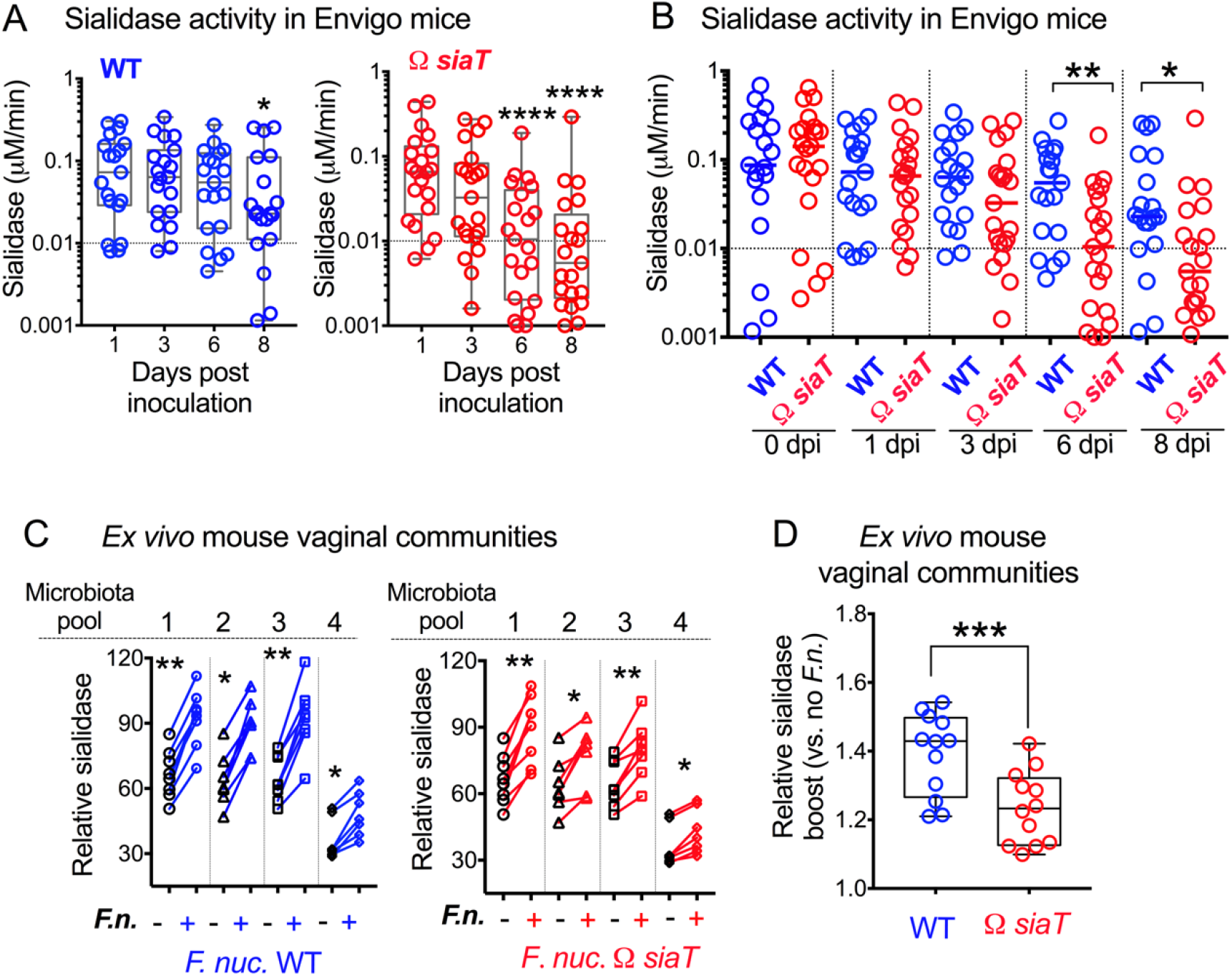
Sialidase activity in *ex vivo* cultured vaginal microbial communities from Envigo mice increases in presence of *F. nucleatum*. (A) Sialidase activity in vaginal washes from 1 to 8 dpi from individual animals purchased from Envigo, estrogenized, and inoculated with either WT or *ΩsiaT F. nucleatum*. Data at later time points were compared to day 1 values using Friedman test, along with correction for multiple planned comparisons using Dunn’s test. (B) Same experiment and data as shown in **A** but analyzed to compare between WT or *ΩsiaT-*inoculated animals at each time point using the Mann-Whitney test. Data in A and B represent combined data from two independent experiments. Data points with negative values were set to 0.001 to represent them on the log scale. (C) Sialidase activity in microbiota pools from Envigo mice. Communities were cultured in the presence or absence of *F. nucleatum* (*F. nuc.*) WT or *ΩsiaT*. Each “microbiota pool” consists of a cultured vaginal community from pooled vaginal wash of 4-5 co-housed mice. Wilcoxon paired-sign rank test was used for pairwise comparison of sialidase activity in each cultured microbiome compared to the identical microbiome cultured in the presence of *F. nucleatum*. Data shown is combined from 4 independent biological replicates with 7-8 technical replicates for each microbiota pool. (D) Same experimental data as in **C**. Data shown is combined from all microbiota pools for each group and analyzed to allow for comparison of the relative sialidase boost between no added *F. nucleatum* (*F.n.*) versus addition of WT or *ΩsiaT*. A statistical comparison between the two groups was performed using Wilcoxon matched-pairs signed rank test. Data shown is combined from 3 independent biological replicates. Line in the bar indicates mean value. On all graphs *P<0.05, **P<0.01, ***P<0.001, ****P<0.0001. See also Figure S6.

To formally test whether *F. nucleatum* could escalate the sialidase activity in vaginal communities, we revived previously cultured and frozen microbiota pools (**Figure 3C**) from vaginal washes of the Envigo mice, which were collected and pooled before *F. nucleatum* inoculation (**Schematic S1, Step 1**). To resurrect each microbiota pool, we gently scraped bacterial communities off the entire solid media plate, then re-suspended and OD-normalized them in supplemented Columbia broth (**Schematic S1, Step 2**). Sialidase activity was measured in the microbiota pools cultured O/N with either media alone (mock) or media containing WT or *ΩsiaT F. nucleatum*. In cultures from each of the four microbiota pools, those that received either WT or *ΩsiaT* had more sialidase activity than the corresponding pools incubated with media alone (**Figure 5C**). However, consistent with the *in vivo* data shown in **Figure 5A-B**, the relative boost in sialidase activity *in vitro* was significantly higher in communities cultured with WT as compared to *ΩsiaT* (**Figure 5D**). As expected, *F. nucleatum* addition did not lead to any increase in sialidase activity in vaginal communities from Jackson (sialidase-negative) mice (**Figure S6**). Thus, while both *F. nucleatum* strains stimulated microbiota-derived sialidase activity, this effect was most apparent with *F. nucleatum* that could take up and consume liberated sialic acids.

### *F. nucleatum* enhances sialidase production by human vaginal communities and facilitates outgrowth of sialidase-positive *Gardnerella*

Our findings in mice led us to wonder whether *F. nucleatum* also escalates the production of sialidase activity in human vaginal microbial communities. To address this question, we collected vaginal swabs, anaerobically transported and eluted vaginal communities into media, and then froze them immediately with cryoprotectant (**Schematic S2, Step 1**). Next, vaginal communities from 21 women, who had sialidase activity in their aerobic vaginal swab eluates (data not shown), were thawed and cultured anaerobically with or without *F. nucleatum* (**Schematic S2, Step 2**). Samples in which *F. nucleatum* was added had significantly higher amounts of sialidase activity than the identical corresponding community cultured without *F. nucleatum* (**Figure 6A**).

**Figure 6.**
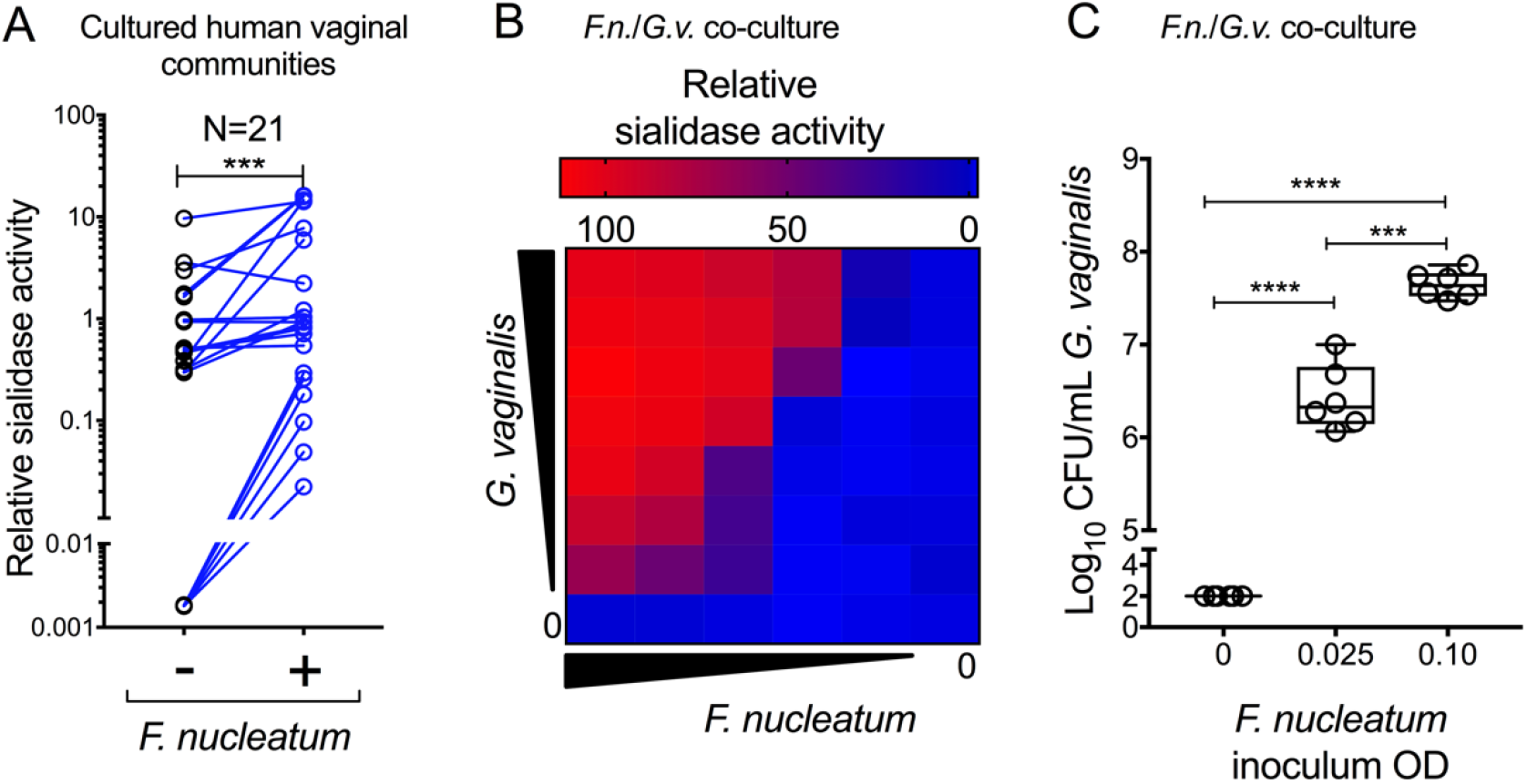
*F. nucleatum* supports growth and sialidase production by human BV bacteria. (A) Human vaginal communities were cultivated anaerobically in Columbia media in the presence or absence of added *F. nucleatum*. Sialidase activity was measured following anaerobic culture. Communities from 21 individual women were used. Data are combined from 2 independent experiments. A statistical comparison between the two groups was performed using Wilcoxon matched-pairs signed rank test. Negative values were set to 0.0018 (lowest positive value) to depict them on the log scale. (B-C) *G. vaginalis* (*G.v.*) was co-cultivated anaerobically in Columbia media in the presence or absence of *F. nucleatum* (*F.n.*), followed by measurement of sialidase activity (B) and viable titers of *G. vaginalis* (colony forming units, C). Note that *G. vaginalis* was not detectable under these conditions in the absence of *F. nucleatum.* In this case, *G. vaginalis* levels were plotted at one half the limit of detection (LOD=200 CFU/mL). Heat map data is representative of two independent experiments. CFU data is combined from two independent experiments, each with 3 technical replicates each. On all graphs ***P<0.001, ****P<0.0001.

There are at least two possible explanations for the enhanced sialidase activity observed upon addition of *F. nucleatum* to vaginal communities. First, if *F. nucleatum* drives sialidase activity in communities by promoting the growth of sialidase-producing bacteria, it would indicate that the organism engages in a mutualistic relationship with sialidase-producing bacteria. On the other hand, *F. nucleatum* could promote expression of sialidase without affecting the growth of sialidase-producing bacteria. In the absence of growth promotion, it would suggest a commensal or parasitic relationship between *F. nucleatum* and sialidase-producing bacteria (depending on whether the latter were unaffected or harmed in the process). To investigate this question, we asked whether *F. nucleatum* had growth-promoting effects on *G. vaginalis*, a bacterium thought to be a primary source of sialidase activity in vaginal fluids of women with BV (Lewis et al., 2013; Lewis et al., 2012). Varying concentrations of *G. vaginalis* (strain JCP8151B) and *F. nucleatum* were co-inoculated into supplemented Columbia broth, which is nutrient rich, but not a preferred media type for *G. vaginalis*. Cultures in 96-well plates were incubated at 37°C, O/N, in the anaerobic chamber. Under these conditions, *G. vaginalis* on its own did not grow or produce detectable sialidase activity. However, addition of *F*. *nucleatum* resulted in statistically significant, dose-dependent increases in both sialidase activity and *G*. *vaginalis* titers (**Figure 6B and 6C**). Remarkably, *F. nucleatum* boosted *G. vaginalis* growth by more than four orders of magnitude under these conditions. Even at the highest inoculum of *G. vaginalis*, sialidase activity was detectable only if *F. nucleatum* was present, which was true even at the lowest inoculum. Taken together with our previous experiments, these data strongly support the interpretation that sialidase-producing vaginal bacteria and *F. nucleatum* mutually benefit each other.

## Discussion

Microorganisms within a biome compete for resources, and their ability to colonize an already occupied niche depends on the ability to access nutrients for growth. We show that *one* resource in particular -- sialic acid -- only becomes accessible to *F. nucleatum* in the presence of sialidases and the ability to utilize this resource encourages its colonization within context of a sialidase-positive microenvironment. We also observed that a mutually beneficial relationship exists between *F. nucleatum* and sialidase producing bacteria in vaginal microbiomes- in both the mouse vagina and in cultures of native vaginal bacterial communities from mouse, as well as humans (despite the fact that these two have very different vaginal microbiotas). Our results show that *F. nucleatum* promotes the growth of sialidase-producing bacteria, such as *G. vaginalis*, often found in abundance within the vaginal microbiome of women with BV. Thus, the data presented here supports a paradigm that microbial sialic acid catabolism is a key metabolic pathway that can shape the vaginal microbiome.

While an ever-growing number of literature reports describe the contents of the human vaginal microbiome, we still know surprisingly little about *how* vaginal dysbiosis develops or why women with BV are more likely to harbor potentially pathogenic vaginal bacteria. Unfortunately, the advancement of mechanistic research on the vaginal microbiota has been hampered by a lack of experimental models. In particular, *in vivo* models have not been developed to study whether and how interactions between vaginal bacteria promote dysbiosis of the vaginal microbiome. Bacteria associated with vaginal dysbiosis are often members of understudied taxonomic groups; they are fastidious and anaerobic, and genetic systems are generally not available. In the present study, we have brought new genetic manipulation, culture-based methods, and an *in vivo* small animal model to bear on these challenges, using *F. nucleatum* as a model organism. By deploying novel experimental systems, we suggest that the ability of potential pathogens to become established in the vagina during dysbiosis is not simply due to the absence of “healthy” lactobacilli and their antimicrobial functions, but also because of mutualistic interrelationships that support the entrance of these new pathogenic members into the community.

*F. nucleatum* often co-inhabits with sialidase producing bacteria at other mucosal niches, such as the oral cavity, gut, and airway. Thus, our findings also have implications in other infections caused by this bacterium. *F. nucleatum* is a key member of the multi-species biofilm seen in periodontal disease, and other members of this biofilm are known sialidase-producers (Fukui et al., 1971; Li et al., 2012; Nonaka et al., 1983). Notably, similar to BV (Lewis et al., 2013), depletion of sialic acids from the oral mucosa has been observed in gingivitis (Davis and Gibbons, 1990), a condition associated with overgrowth of *F. nucleatum*. Recently, *F. nucleatum* has been identified as a cause of colorectal cancer (Brennan and Garrett, 2016). Moreover, colorectal tumors that have metastasized to other locations in the body commonly contain both *F. nucleatum* and *Bacteroides* (known sialidase producers) (Bullman et al., 2017). In fact, sialidase-producing *Bacteroidetes* are often found side by side with *Fusobacteria* in abscesses, amniotic fluid infections, and polymicrobial bacteremia/sepsis (Denes and Barraud, 2016; Sakamoto et al., 1998; Siqueira et al., 2001; Song et al., 2014; Urushiyama et al., 2017). Together with the data presented here, these findings suggest that *F. nucleatum* may benefit from sialidase-producing bacteria during disease processes throughout the human body.

In conclusion, we provide a unique example in which sialic acid catabolism forges a symbiotic relationship between microbes, which in turn encourages and allows the maintenance of a (sialidase-producing) dysbiotic microbial community in the reproductive tract. These findings advance our understanding of microbial interactions that promote vaginal dysbiosis and may help explain why women with BV are more susceptible to reproductive tract infection.

## Acknowledgements

We thank Nadum Member-Meneh and Courtney Amegashie for technical contributions to clinical sample collection and processing, Justin Fay and Andrew Kau for advice on microbiome profiling, Justin Merritt for providing some of the *Fusobacterium* strains, Deborah Frank for editorial assistance and advice, and Marcy Hartstein for assistance with illustrations. We also thank Denise Spear WHNP-BC, and Valerie Higginbotham WHNP-BC, for their participation in collecting vaginal swabs from participants and the St. Louis County Department of Public Health for facilitating this study. Finally, we thank the women who have generously provided their informed consent and vaginal samples for use in these studies.

## Author Contributions

Study design, A.L. and W.L.; Writing of the first draft of the manuscript, K.A., A.L., W.L.; Editing and final approval of the manuscript, all co-authors; Data analysis and interpretation, K.A., A.L., W.L., L.R., N.G. and B.T.; Bioinformatics, K.A., B.T., W.L.; Cloning and mutagenesis: L.R. and H.L.; *In vitro s*ialic acid foraging and growth experiments: L.R., J.P., W.L., H.L. and K.A.; Mouse handling and experimentation: K.A., L.F. and V.P.O.; 16s library preparation and sequencing: B.T.; Microbiome data analysis: B.T., W.L., K.A. and A.L.; management of human study data and sample collection process: H.R., A.L. and W.L.; Design and execution of mouse and human *ex-vivo* community experiments: K.A. and W.L.

## STAR * Methods

### KEY RESOURCES TABLE

Bacterial strains, plasmids and primers (see **Table S1**)

## METHOD DETAILS

### Bioinformatic analysis

Bioinformatics analysis was conducted using protein blast (BLASTp ver. 2.2.31+) program from the BLAST suite (Altschul et al., 1997). Amino acid sequences of *N*-acetylneuraminate lyase (NanA) from *E. coli* MG1655 (GenBank: AAC76257.1) and *F. nucleatum* subsp. *nucleatum* ATCC 23726 (GenBank: EFG95907.1) were used as query to non-redundant (NR) protein sequences database for *F. nucleatum* (taxid: 851). Any hits with an *E*-value less than 1e-9 were considered significant.

### Bacterial strains and growth conditions

Wild-type (WT) strains of *E. coli* [TOP10, Invitrogen and MG1655, UTI89 and CFT073 (provided by Scott Hultgren)] were grown in lysogeny broth (LB) at 37°C unless otherwise noted. For plasmid maintenance, ampicillin was used at 100 μg/mL and chloramphenicol at 20 μg/mL. *Gardnerella vaginalis* JCP8151B (Lewis et al., 2016; Lewis et al., 2013), hereafter simply referred to simply as *G. vaginalis*, was cultured in NYC-III media containing 10% heat inactivated horse serum at 37°C in an anaerobic chamber. *Fusobacterium nucleatum* strains ATCC25586, ATCC23726 and ATCC10953 were obtained from ATCC. Strains JMP2A, FQG51A, JMSY1, JMH52, SYJL4, JM6, and JM4 (Yoneda et al., 2014) were provided by Justin Merritt. ATCC23726 is the primary strain used to model interaction with sialidases and the microbiome and is hereafter referred to simply as *F. nucleatum*. All *Fusobacterium* strains were grown at 37°C in an anaerobic chamber either on Columbia blood agar plates (containing 5% defibrinated laked sheep blood) or liquid Columbia media, both supplemented with Hemin (5 μ g/mL) and Vitamin K3 (1 μ g/mL), referred to in the text as “supplemented Columbia” (unless otherwise indicated). *Prevotella bivia* was grown in CDC media supplemented with laked sheep blood (5%), Vitamin K1 (10 μ g/mL) and Hemin (5 μ g/mL) under anaerobic conditions.

### Isolation of a streptomycin-resistant mutant of *F. nucleatum*

*F. nucleatum* ATCC23726 was cultured anaerobically overnight (O/N) in Columbia media at 37°C. Cultured bacteria (20 mL) were pelleted by centrifugation at 16,000 x g for 10 min at room temperature, re-suspended in 1 mL PBS, and plated on Columbia blood plates with 1 mg/mL streptomycin sulfate to select for spontaneous streptomycin-resistant (SmR) colonies. After two days, streptomycin-resistant colonies from Columbia plates (with strep) were cultured in liquid Columbia media and used for subsequent experiments. Mutation of *rpsL* in the *F. nucleatum* 23726-SmR strain was confirmed by colony PCR and DNA sequencing.

### Mutagenesis targeting *siaT* in *F. nucleatum* ATCC23726

Restriction endonucleases, Phusion DNA polymerase, and T4 ligase were obtained from New England Biolabs. First, a suicide vector was constructed using the origin of replication from pUC19 and the chloramphenicol acetyltransferase gene (*catP*) from pJIR418 (see **Table S1** for more info on plasmids and primers). Briefly, the origin of replication was amplified from pUC19 using primers pUC19 ori F Sac Kpn Nco and pUC19 ori R Bgl2. The *catP* gene from pJIR418 was amplified with primers catP F Bgl2 and catP R Bam Pst Nco. The two amplicons were desalted, digested with NcoI and BglII, and ligated together, resulting in pLR23. A 0.5 kb internal fragment of *F. nucleatum siaT* was amplified from ATCC23726 chromosomal DNA with the primers 23726 nan F Nco and 23726 nanR Sac. Following desalting, this PCR product and pLR23 were digested with NcoI and SacI and ligated together, creating pLR25. The streptomycin-resistant strain of *F. nucleatum* ATCC23726 was used as the background for the *siaT* mutation. A starter culture of this strain was diluted 1:1000 into 10 mL Columbia broth and grown O/N in the anaerobic chamber at 37°C. Bacteria were then pelleted at 12,000 x g for 10 min and washed twice in 1 ml of 1 mM MgCl_2_ containing 10% glycerol. The bacteria were washed twice more in 1 mM MOPS with 20% glycerol, and resuspended in 100 μl of the same buffer. Next, 20 μ g of purified pLR25 was mixed with the above suspension and incubated on ice for 10 minutes. The suspension was then transferred to a cuvette with a 0.1 cm gap and electroporated in an Eppendorf Electroporator 2510 set to 2.5 kV. Bacteria were recovered anaerobically for 5 hours in Columbia broth supplemented with 1 mM MgCl_2_ at 37°C, then plated on Columbia agar with 5 μ g/mL thiamphenicol. Plasmid integration at the desired locus was confirmed by colony PCR using the primers 23726 nan test F1 and pLR24 5’ F (**Figure S2**). The mutant strain is referred to as *F. nucleatum ΩsiaT*.

### Complementation of *E. coli* MG1655 *ΔnanA* and growth comparison

MG1655 *nanA* was amplified with the primers MG1655 nanA F Nco and MG1655 nanA R Pst. The PCR product was desalted, digested with the restriction enzymes NcoI and PstI, and ligated into the NcoI and PstI sites of pTrc99A to create pLR7. *F. nucleatum* ATCC 23726 *nanA* was amplified with the primers Fuso nanA F Nco and Fuso nanA his R Bam. Following desalting and digestion with NcoI and BamHI, the amplicon was cloned into the same sites in pTrc99A, creating pLR10. The *E. coli* MG1655 *ΔnanA* mutant LSR4 was described previously (Robinson et al., 2017). LSR4 was transformed with pLR7, pLR10, or the vector control (pTrc99A) (see **Table S1**). The three strains were grown to an optical density of 1.0 in LB broth containing 100 μ g/mL ampicillin. Then, each strain was diluted 1:100 into M63 media lacking glycerol and supplemented with 15 mM Neu5Ac. Optical density at 600 nm was measured every 10 minutes at 37°C in 96 well plate (flat bottom, Greiner) in a TECAN M200 plate reader with 10 seconds of orbital shaking at the beginning of each time point.

### Sialate lyase assay

The *E. coli ΔnanA* mutant (LSR4) containing pLR7 (*E. coli* MG1655 *nanA*), pLR10 (*F. nucleatum nanA*), or vector control (pTrc99A) was grown shaking at 37°C in 35 mL Circle Grow broth supplemented with 100 μ g/mL ampicillin to an optical density of 1.0. Cultures were then transferred to room temperature, induced with 0.2 mM IPTG, and grown shaking O/N. By the next morning, cultures had grown to an optical density of approximately 4.0. Next, 25 ml of each culture was centrifuged at 12,000 x g for 10 minutes and the cells were resuspended in 800 μl PBS. Cell suspensions were sonicated on ice for 40 seconds in 1 second bursts at 24% amplitude in a Sonic Dismembrator (Fisher Scientific). Insoluble debris was removed by centrifugation at 15,000 x g for 10 min, and 90 μL of each supernatant was mixed with 10 μL of 1 mM Neu5Ac for a final concentration of 100 μM. Mixtures were incubated at 37°C and at each time point 10 μL samples were removed, diluted 5x in water, and frozen at −20°C prior to analysis by DMB-HPLC.

### Measurement of total and free sialic acids by DMB-HPLC

Derivatization and quantitation of sialic acids by HPLC was carried out as previously described (Lewis et al., 2013). Briefly, total sialic acids were measured by first releasing any bound sialic acids using mild acetic acid hydrolysis (2 N acetic acid for 3 h at 80°C), followed by derivatization of all free sialic acids with DMB (1,2-diamino-4,5-methylenedioxybenzene) as described below. Alternatively, free sialic acid levels were measured by DMB derivatization without prior acid hydrolysis. Thus, the concentration of bound sialic acids is measured indirectly (bound = total-free). Reaction conditions for DMB derivatization were 7 mM DMB, 22 mM sodium thiosulfite, 0.75 M 2-mercaptoethanol, and 1.4 M acetic acid for 2 h at 50°C. Derivatized samples were injected into a Waters HPLC equipped with a reverse-phase C18 column (Tosoh Bioscience) and eluted using isocratic conditions at 0.9 ml/min using 8% methanol, 7% acetonitrile in water. An on-line fluorescence detector (Waters) was set to excite at 373 nm and detect emission at 448 nm. Peak integrations were used to quantitate sialic acid content by referencing a standard curve of pure sialic acid (Neu5Ac, Sigma) derivatized in parallel.

### Free sialic acid foraging by *F. nucleatum*

*E. coli* (Top 10) was cultured aerobically and *F. nucleatum* strains (see **Table S1**) were cultured anaerobically in Columbia media supplemented with 100 μM sialic acid (Neu5Ac, Sigma, A0812). After 24 - 48 h of growth, media supernatant was separated from bacterial cells by centrifugation at 15,000 x g. In some experiments, collected media supernatant was filtered using a 10K molecular weight cut off (MWCO) filter. Samples were then processed for DMB derivatization for measurement of free sialic acid (Neu5Ac) remaining in the medium as described above.

### Sialidase prepared using protease and affinity chromatography

One liter of *G. vaginalis* was grown O/N to an OD_600_ of 1.6. Cells were pelleted at ~18,000 x g and washed with 60 mL of 0.1M sodium acetate buffer pH 5.5, centrifuged again, then re-suspended in 20 mL sodium acetate buffer. Freshly dissolved subtilisin (Sigma) was added to the well-mixed cell suspension at a final concentration of 50 μ g/mL and rotated at 37°C for 2 hours. Bacteria were pelleted at 15,000 x g and the supernatant was collected. A fresh batch of 200 mM PMSF in ethanol was prepared, 100 μL was added to the supernatant for 1 mM final, mixed for 15 minutes and then stored at 4°C O/N. After brief centrifugation and sterile filtration through a 0.4 micron membrane, 250 μL of the preparation was applied to an affinity column to enrich for sialidase. Specifically, well-mixed phenyl oxamate agarose beads (Sigma), were applied as a 1 mL slurry to a fritted polypropylene column and equilibrated with sodium acetate buffer to give a packed bed of ~0.5 cm^2^. 100 μL of subtilisin-released material was applied to the affinity column and washed with 5 x 0.7 mL sodium acetate buffer. Fractions of 250 μL were collected in a polypropylene plate during washing. Elution was carried out with a total of 4.5mL 100 mM NaHCO_3_ pH 9.5, and 150 μL fractions were collected. Sialidase activity was monitored in small samples of column fractions by adaptation of the BV blue kit (Gryphus) to a 96-well microtiter plate assay as follows: 5 μL aliquots of column eluate were combined with 20 μL of BV blue reagent solution in a light-colored V-bottom or U-bottom 96-well plate on a white background. After 10 minutes, one drop of developer solution was added to the reaction mixture in each well. Fractions with sialidase activity were identified by the blue color in the corresponding BV blue reaction, and the sialidase positive fractions were pooled, then concentrated to approximately 50 μL using a molecular weight cutoff filter (Amicon).

### Foraging of sialic acids from bound sialoglycans in culture media by *F. nucleatum* in the presence or absence of exogenous sialidases

These experiments were performed in two different media types (Columbia media and CDC media) and each condition was performed twice independently. Each type of media was treated O/N with sialidases from different sources. As a positive control, purified recombinant *Arthrobacter ureafaciens* sialidase (AUS, EY Laboratories, Inc.), resuspended in 100 mM sodium acetate at 5 mU/μL, was used to treat media by adding 1 μL enzyme for each 499 μL of culture media. A parallel batch of media was mock treated with 100 mM sodium acetate (also 1:500). Additionally, media was treated with *G. vaginalis* sialidase, released and purified from *Gardnerella* cultures as described above (added to media at 1:500). In the experiments performed in CDC media, we also used an additional sialidase-producing bacterium associated with BV -- *Prevotella bivia*. For the latter, cultured bacteria in CDC media were pelleted and lysed in water. Then, 50 μL of this crude lysate was added to each 450 μl of CDC media. Each of the media conditions was allowed to incubate O/N in the anaerobic chamber, followed by inoculation of *F. nucleatum* strains ATCC23726 or ATCC25586 the following day. The zero time point samples were taken immediately after inoculation by removing an aliquot of media. After culture of the remaining sample for 48 h, another aliquot was removed. Samples taken at the zero and 48 h time points were subjected to DMB-HPLC as described above. The following formulas were used to calculate the values shown:

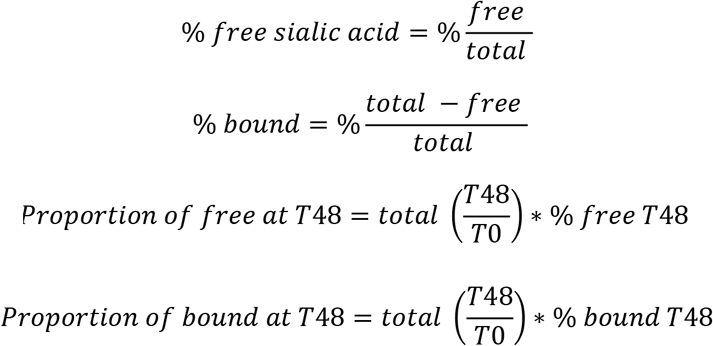

### Fluorometric assay for sialidase activity

All sialidase assays were performed in 96 well U-bottom black polypropylene plates (Eppendorf). Plates were sealed with an optical clear film and substrate hydrolysis was monitored by measuring fluorescence of 4-methylumbelliferone (4-MU) (Excitation 365 nm, Emission 440 nm) in a Tecan M200 plate reader every 2 min for 1-2 h at 37°C. To test for potential sialidase activity produced by *G. vaginalis* and *F. nucleatum*, O/N cultures were adjusted to an OD_600_ of 0.5 and 25 μL was mixed with an equal volume of PBS in the plate, followed by addition of 100 μL Dulbecco’s PBS containing 100 μM 4-methylumbelliferyl-Neu5Ac (4MU-sialic acid--Goldbio). To monitor sialidase activity in mouse vaginal washes, 25 μL of vaginal wash was mixed with 50 μL PBS containing 100 μM 4MU-sialic acid. 4MU in PBS (0 – 67 μM), was included in sialidase assay with mouse vaginal washes to calculate the rate of 4MU-sialic acid hydrolysis by comparison to standards. Rate of 4MU-sialic acid hydrolysis was used to compare sialidase activity in the mouse vaginal washes between different time points or over a period of time. For *in vitro* experiments examining the effect of *F. nucleatum* on cultured mouse/human vaginal specimens, or *G. vaginalis*, 50 μL of the final culture was mixed with 100 μL sodium acetate buffer (pH 5.5) containing 500 μM 4MU-sialic acid. Similarly, for sialidase activity assays on (uncultured) human vaginal specimens, 50 μL of re-suspended swab eluates was transferred to the plate and mixed with 100 μL sodium acetate buffer (pH 5.5) containing 500 μM 4MU-sialic acid. 4-MU in sodium acetate buffer (0-300 μM), was used as a standard in sialidase assay with vaginal swab eluates.

### Growth comparison of *F. nucleatum* WT and *ΩsiaT*

*F. nucleatum* wild-type and *ΩsiaT* were grown O/N in NYCIII media, then adjusted to reach a final starting OD_600_ of 0.05 in fresh media containing different carbohydrate sources as follows: (i) NYCIII media with 24.4 mM glucose, (ii) NYCIII media with no glucose (low carbohydrate) and (iii) NYCIII no glucose media supplemented with 24.4 mM Neu5Ac. Media alone or inoculated with *F. nucleatum* was transferred to a clear 96 well plate (100 μL/well), followed by overlaying 50 μL mineral oil and sealing the plate with a transparent seal to maintain anaerobic conditions. Growth was monitored by recording OD at 600 nm every 15 min at 37°C in a TECAN plate reader for ~45 h.

### Ethics statement

Mouse experiments were carried out in strict accordance with the recommendations in the Guide for the Care and Use of Laboratory Animals. The protocol was approved by the Animal Studies Committee of Washington University School of Medicine (Protocol Number: 20110149).

### Mouse vendor information, housing conditions and general handling procedures

For experiments with *F. nucleatum*, female C57BL/6 mice (6-7 weeks old) were obtained from Jackson Labs (Bar Harbor facility) and Envigo (202-A Indianapolis facility) and were received between Jan. 2016 and Jan. 2017. For experiments testing vaginal sialidase activity among different mouse vendors, female C57BL/6 mice were also obtained from Charles River (NCI Grantee) and Taconic. Once received at Washington University, mice were housed in a barrier facility at 70 +/- 2°F, 12:12 light:dark cycle, corncob bedding, and nestlets for enrichment changed at least once per week. Mice were allowed to rest for approximately 1 week prior to the commencement of procedures. Mice were estrogenized as indicated in figure legends by intraperitoneal injection under isofluorane anesthesia with 0.5 mg β-estradiol in 100 μL filter-sterilized sesame oil (Sigma). Vaginal washes were collected from isofluorane anaesthetized animals by gently inserting a pipette tip with 50 μL of sterile Dulbecco’s Phosphate Buffered Saline ~2-5 mm into the vagina and pipetting up and down.

### Mouse model of vaginal colonization with *F. nucleatum*

To distinguish inoculated *F. nucleatum* from endogenous members of the microbiota for enumeration of colony forming units, we used a spontaneous streptomycin-resistant isolate derived from ATCC 23726, which we refer to simply as “wild-type” (WT) *F. nucleatum*. The *ΩsiaT* strain described earlier was derived from this WT strain. Mice were treated with β- estradiol (see above) three days prior to, and on the day of, inoculation with all animals receiving the first dose ~1 week post arrival. A total of 80 mice (40 mice from each vendor) were used in this study. Two independent experiments of *F. nucleatum* colonization were conducted approximately 1 year apart with each individual experiment including 20 mice from Envigo labs and 20 from Jackson labs. Ten mice per vendor were vaginally colonized with *F. nucleatum* WT and 10 with the *F. nucleatum ΩsiaT*. To prepare the *F. nucleatum* inoculum, the O/N culture was adjusted to an OD_600_ of 4.0 in Columbia media and aliquoted in 1.5 mL eppendorf tubes (25 μL/ tube) for each individual mouse. Mice were vaginally inoculated with ~10^8^ CFU of streptomycin resistant *F. nucleatum* ATCC23726 wild-type or *ΩsiaT* mutant in 20 μL Columbia media using a separate aliquot of inoculum for each mouse. Vaginal washes were collected one day before estradiol was first administered, after estrogenization (on the day of inoculation) and then 3 times per week (on alternate days) until 38 days post inoculation (dpi). Mice were considered “cleared” of *F. nucleatum* when two consecutive washes were negative for *F. nucleatum* CFU. Once cleared, mice did not re-acquire *F. nucleatum*, even if co-housed with colonized mice. Washes were collected from all mice by flushing vaginas with 50 μL sterile PBS and rinsing into an additional 20 μL PBS in a sterile 1.5 mL Eppendorf tube. All washes were collected (from 40 mice) within 1 hour and cycled into the anaerobic chamber for colony forming units (CFU) enumeration. *F. nucleatum* titers were determined from washes by preparing 10-fold serial dilutions in PBS and spotting 5 μL of each dilution in quadruplicate onto 1 mg/mL streptomycin selection plates (Columbia plates with defibrinated sheep blood). After anaerobic incubation at 37°C, colonies were then enumerated and reported as recovered CFU per mL of vaginal fluid. Sialidase activity was measured in freshly-collected mouse vaginal washes as described above.

### Mouse model of vaginal inoculation with *E. coli*

Female C57BL/6 mice, 5-7 week old, were obtained from the Envigo (202-A Indianapolis facility). At 48 and 24 hours prior to infection, mice underwent intraperitoneal injections of 0.5 mg β-estradiol 17-valerate (Sigma) in 100 μl filter-sterilized sesame oil (Fisher) to synchronize the mice in estrus. Mice were infected with model uropathogenic *Escherichia coli* strains (UTI89 and CFT073 (Mobley et al., 1990; Mulvey et al., 2001) or mutant derivatives thereof. Strains were cultured statically in lysogeny broth (LB) at 37°C for two consecutive O/N passages. At time of infection, 10^4^ CFU of each *E. coli* strain were inoculated into the vaginas in 20 μl volumes of PBS. To monitor infection and sialidase status, vaginal washes were collected as described above, serially diluted and plated on MacConkey agar (titers of these uropathogenic strains went up to ~10^8^ cfu by 24 hpi). Sialidase activity was measured in freshly-collected mouse vaginal washes as described above.

### *Ex-vivo* mouse vaginal community cultures

Mouse vaginal wash material remaining after sialidase assays and *F. nucleatum* CFU analysis was pooled by cage (5 mice per cage) for each vendor or treatment condition. A small portion (5 μL) of each pooled vaginal wash sample was used as inoculum to culture the vaginal bacterial communities in supplemented Columbia broth under anaerobic conditions. We refer to these vaginal communities as “microbiota pools”. Each microbiota pool represents either uncultured pooled vaginal wash or cultured bacterial community from pooled vaginal wash of mice co-housed in the same cage. The cultured communities (1^st^ passage) were frozen with 20% glycerol after O/N growth. Remaining pooled vaginal washes were stored at −20°C for later microbiome studies (see below). See also Schematic S1.

### DNA extraction and community profiling

For microbiome analysis by 16s sequencing, pooled vaginal washes (uncultured microbiotas) from 8 different cages (4 for each vendor, 1 specimen per cage) were centrifuged at 16,000 x g for 5 min, supernatant was removed, and the pellet was used for DNA extraction. For cultured vaginal communities (4 for each vendor, 1 specimen per cage), communities frozen in supplemented Columbia broth were streaked out on Columbia blood plates and grown O/N at 37°C in anaerobic chamber (2^nd^ passage). Colonies from Columbia blood plates were gently scraped off using a cell scraper and re-suspended in liquid Columbia media. For DNA extraction, bacterial cells from these communities were separated from the supernatant by centrifugation at 16,000 x g for 5 min. DNA was extracted using the Wizard Genomic DNA Purification kit from Promega (Cat. No. A1120) following supplier instructions. See also Schematic S1.

### Mouse vaginal community analysis by V1-V2 sequencing of the 16S rRNA gene

We amplified the V1-V2 region of the 16s ribosomal subunit with the universal 27F and 338R primers (Ravel et al., 2011). Both the primers contained a common adaptor sequence and the forward primer also contained a barcode sequence for multiplexing. The primers were as follows: 338R-5’-AGACGTGTGCTCTTCCGATCTCAT**GCTGCCTCCCGTAGGAGT**-3’ and 27F-5’-ACGACGCTCTTCCGATCTNNNNNNNNCT**AGAGTTTGATCCTGGCTCAG**-3’. Where the bolded sequences denote the universal primers and 8-bp barcode is denoted by 8 Ns. After the V1-V2 region was amplified using the 27F and 338R primer pair, the PCR product was treated with Exo-SAP-IT to remove primers. A second PCR was performed to add unique indexes for further multiplexing. The subsequent amplicons were quantified and pooled. The pool was run on a 0.8% agarose gel, excised, and extracted. Agencourt AMPure XP beads were used for further size selection and purification. 2 x 150 paired-end sequencing was completed using the Illumina MiniSeq platform through the Center for Genome Science at Washington University School of Medicine. Reads were trimmed for adaptor sequences using ea-utils fastq-mcf version 1.04.676 and de-multiplexed based on the unique barcode and index identifiers. USEARCH (Edgar, 2010) version 10.0.240 was used for the following USEARCH commands: (i) Reads were merged using the fastq_mergepairs command, (ii) quality filtered with a maximum expected error of 1 and a minimum sequence length of 100 using the fastq_filter command, (iii) merged reads were then dereplicated using the fastx_uniques command, and (iv) OTUs were clustered using the cluster_otus command or unoise3. As part of the cluster_otus and unoise3 commands, chimeric sequences were filtered out and discarded and a minimum abundance of 2 reads per OTU was applied. OTUs were assigned taxonomic predictions using the RDP 16s database (version 16) with a confidence threshold of 0.8. Samples were rarified to 1000 reads per sample. All OTUs at >1% abundance were included in analysis.

### *F. nucleatum*’s influence on sialidase activity in mouse vaginal communities

Experiments were done with microbial communities cultured using pooled vaginal wash specimens (referred to as “microbiota pools”) collected from both Envigo and Jackson mice after estrogenization, but before inoculation with *F. nucleatum*. Cultured mouse vaginal communities (1^st^ passage) frozen in Columbia media were streaked out on Columbia blood plates and grown in an anaerobic chamber at 37°C (2^nd^ passage). After two days, Columbia blood plates were gently scraped using an inoculating loop and bacteria were re-suspended in supplemented Columbia broth. The OD_600_ of the suspension was checked and adjusted to 0.5 for the vaginal communities and 1.0 for *F. nucleatum* strains. Experiments were done in v-bottom, 96 well, sterile, deep well (2 mL) plates (Eppendorf). *F. nucleatum* was inoculated in 700 μL (total volume of media per well) of Columbia broth at a starting OD_600_ of either 0.10 or 0.05 or 0.025, followed by addition of vaginal communities at starting OD_600_ 0.05. Appropriate controls, such as uninoculated columbia broth, columbia broth inoculated with *F. nucleatum* alone at a starting OD_600_ of 0.10 and columbia broth inoculated with each individual vaginal microbial community alone were included in each experiment. After the addition of aluminum foil seals (Beckman Coulter, Product ID: BK538619), plates were incubated O/N (approx. 16.5 h) at 37°C in the anaerobic chamber. After O/N growth (3^rd^ passage of mouse communities), sialidase activity was checked in each well using the fluorometric sialidase assay as described earlier in the methods. See also Schematic S1.

### Collection of Human vaginal communities

According to Washington University IRB-approved protocol number 201704121 (PI:Amanda Lewis), we enrolled nonpregnant women seeking care at the North Central Community Health Clinic, a no fee clinic operated by the St. Louis County health department. While no specific racial or demographic group was sought or excluded for participation in this study, >90% of women enrolled self-identified as black, a demographic very similar to the community neighboring the clinic. Women with known Human Immunodeficiency Virus or Hepatitis C Virus or with recent antibiotic use (past 3 weeks) were excluded. Mid-vaginal swabs were collected during speculum exam by a clinician, immediately submerged into pre-reduced Cary Blair media using Starswab Anaerobic Transport System S120D and transported to the laboratory for same day processing. Once in the anaerobic chamber, swabs were eluted in 2X NYCIII media, and “fresh frozen” (i.e. “0 passage,” without growth or amplification of any kind) in the presence of sterile glycerol (20% final). Standard (aerobic) rayon swabs (Starplex double headed, S09D) collected at the same visit were eluted in 1 mL of pH 5.5 sodium acetate buffer and sialidase activity was measured as described earlier in the methods. For co-culture experiments with *F. nucleatum*, we selected anaerobic fresh frozen vaginal microbial communities from women whose aerobic vaginal swab eluates had detectable sialidase activity. Fresh frozen communities from 21 sialidase-positive women were selected. See also Schematic S2.

### *F. nucleatum*’s influence on sialidase activity in Human vaginal communities

*F. nucleatum* ATCC23726 wild-type was streaked out on Columbia blood plates and grown in anaerobic chamber at 37°C for 24 h. The next day, *F. nucleatum* colonies from Columbia blood plates were gently scraped off using an inoculating loop and re-suspended in supplemented Columbia broth. The OD_600_ of the suspension of *F. nucleatum* was adjusted to 1.0. *F. nucleatum* inocula were prepared by 2-fold serial dilution in supplemented Columbia media. Fresh frozen (passage 0) human vaginal communities (N=21) were thawed on ice and diluted 4-fold in 1X NYCIII with no added glucose (low carbohydrate). Experiments were done in v-bottom, 96 well, sterile, deep well (2 mL) plates (Eppendorf). *F. nucleatum* was inoculated at a starting OD_600_ of 0.10, followed by addition of 116 μL of the diluted fresh frozen community in a total of 700 μL media (supplemented Columbia broth) per well. Mock vaginal communities were included as control in each experiment; these mock communities were originally prepared from blank swabs that were eluted and stored using the same media and reagents, in parallel with swabs returning from the clinic. The plates were sealed with a sealing aluminum foil (Beckman Coulter, Product ID: BK538619) and incubated O/N (17 h) at 37°C in the anaerobic chamber. After O/N growth (1^st^ passage of human communities), sialidase activity was measured in each well using the fluorometric sialidase assay as described earlier in the methods. See also Schematic S2.

### *Fusobacterium* influence on growth of *G. vaginalis* and production of sialidase

*F. nucleatum* ATCC23726 wild-type (SmR) and *G. vaginalis* 8151B (SmR) were streaked out on Columbia blood plates and NYCIII plates with streptomycin (1 mg/mL), respectively, and grown in anaerobic chamber at 37°C for 24 h. The next day, colonies from the plates were gently scraped off using an inoculating loop and re-suspended in supplemented Columbia broth. The OD_600_ of the suspension was checked and adjusted to 0.5 for *G. vaginalis* and 1.0 for *F. nucleatum*. A series of inocula were prepared for each strain by 2-fold serial dilution. Experiments were done in v-bottom, 96 well, sterile, deep well (2 mL) plates (Eppendorf). Inocula for both *F. nucleatum* and *G. vaginalis* were then diluted an additional 10-fold in a total of 700 μL of supplemented Columbia broth per well. The plate was sealed with a sealing aluminum foil (Beckman Coulter) and incubated O/N (17 h) at 37°C in the anaerobic chamber. After O/N growth, sialidase activity was measured in each well using the fluorometric sialidase assay as described earlier. *G. vaginalis* titers were determined after O/N growth for a subset of the wells, with and without *Fusobacterium,* at the indicated inocula by preparing 10-fold serial dilutions in PBS (in the anaerobic chamber) and spotting 5 μL of each dilution in quadruplicate onto NYCIII plates containing 1 mg/ml streptomycin, 10 μg/ml colistin, and 10 μg/ml nalidixic acid (colistin and nalidixic acid select against Gram negative bacteria and allow the specific enumeration of *Gardnerella* in this assay).

### Statistical analysis

GraphPad Prism 7.0 software was used for all statistical analyses presented. The statistical tests used to analyze each set of data are indicated in the figure legends. For animal experiments, the figures with titers or sialidase activity show individual data with each data point representing a value from a different animal with a line at the median. For non-parametric analyses, differences between the experimental groups were analyzed with a two-tailed unpaired Mann-Whitney U test. In experiments where data sets are paired, differences were analyzed using Wilcoxon matched-pairs sign rank test. Survival analysis was done using the Gehan-Breslow-Wilcoxon test.

## Supplemental Information

**Schematic S1.**
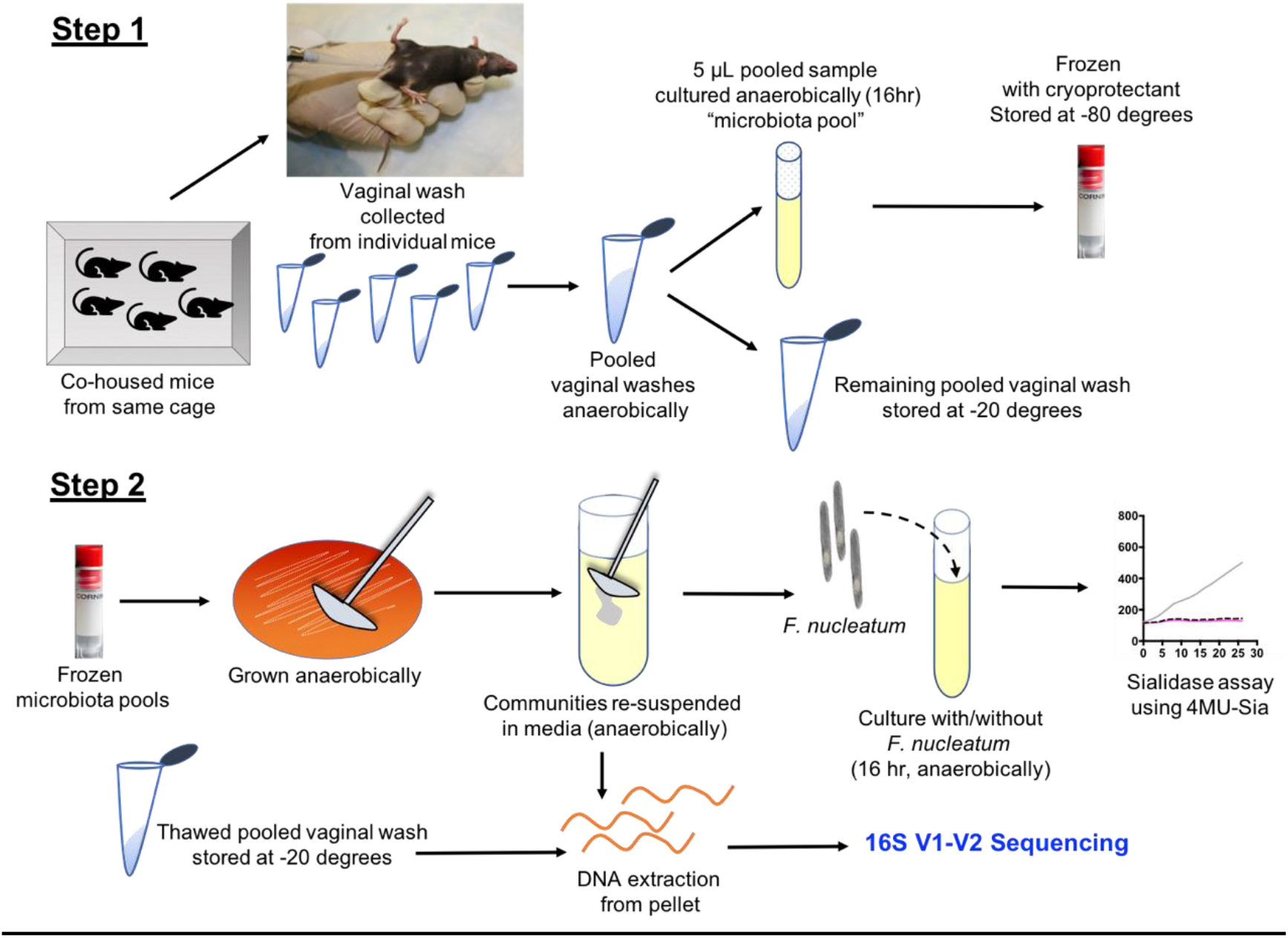
Mouse vaginal microbial communities-collection and amplification. **Step 1** - *Collection of mouse vaginal washes. Washes were pooled from mice co-housed in the same cage. A portion of vaginal wash pools was cultured overnight anaerobically (referred to as “microbiota pools”) to amplify the vaginal bacteria and frozen for subsequent use. Remaining pooled material was stored at −20°C. **Step 2** - On the day of the experiment, frozen microbiota pools were used to recover mouse vaginal bacteria by streaking out on Columbia blood plates in anaerobic chamber and incubating for 24 h at 37°C. Colonies from these plates were re-suspended in liquid media either (a) for DNA extraction, or (b) for co-culture experiments with *F. nucleatum*. *Publication of this animal image was approved by IACUC.

**Schematic S2.**
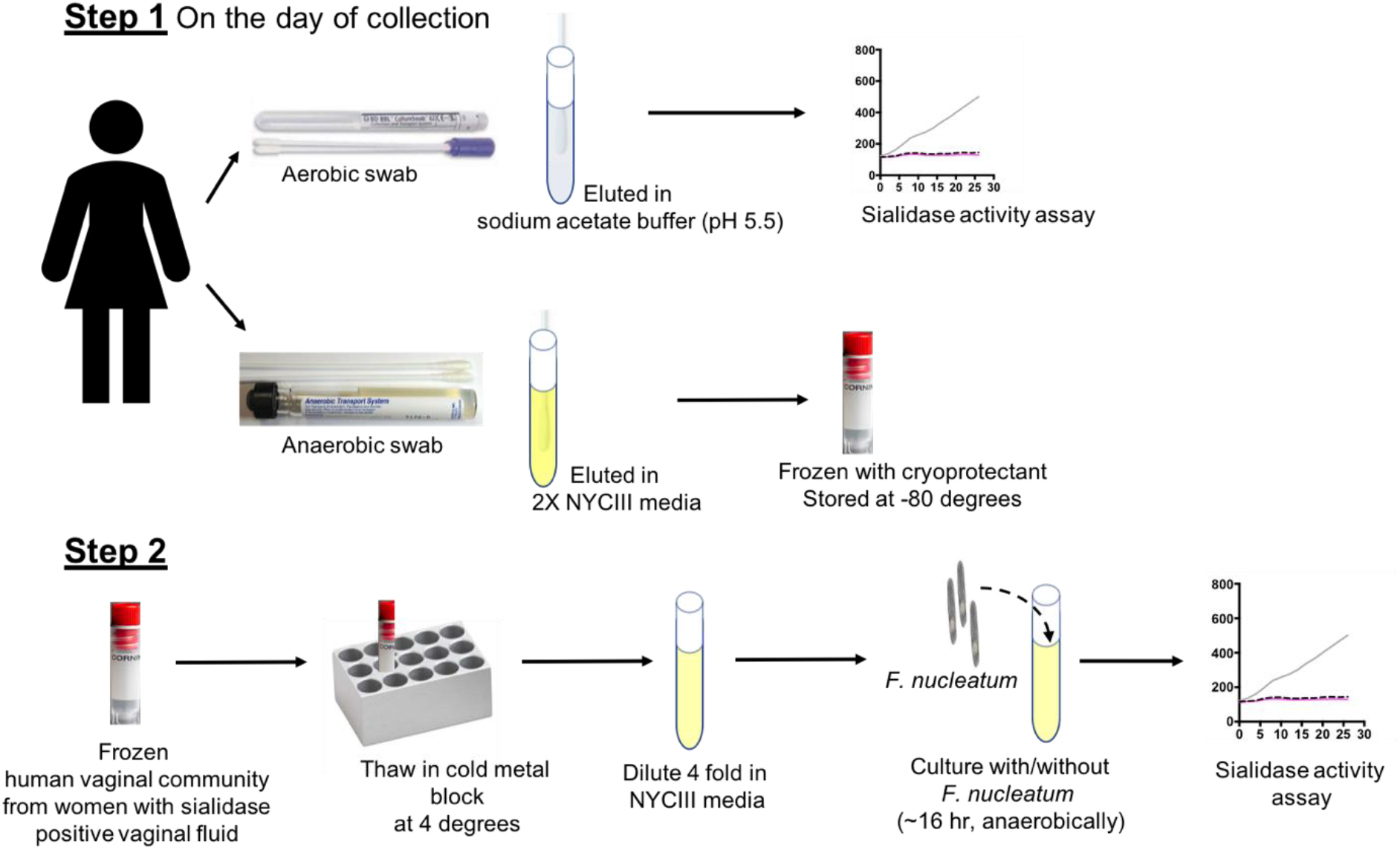
Human vaginal microbial communities-collection and amplification. **Step 1** - Anaerobic and aerobic vaginal swabs were collected on the same day from each participant. Aerobic swabs were eluted in sodium acetate buffer (pH 5.5) and sialidase activity was checked in the swab eluates using fluorogenic 4MU-Neu5Ac substrate. Anaerobic swabs were eluted in 2X NYCIII media (in an anaerobic chamber) and the communities were “fresh frozen”, <u>without any amplification / overnight culture</u>, by mixing with cryoprotectant and storing at −80°C. **Step 2** - On the day of the experiment - fresh frozen anaerobic vaginal communities, from women who had detectable sialidase activity in their aerobic swab eluates, were thawed at 4°C and diluted 4-fold in NYCIII media (in an anaerobic chamber). The diluted communities were used for co-culture experiments with *F. nucleatum*.

**Table S1.**
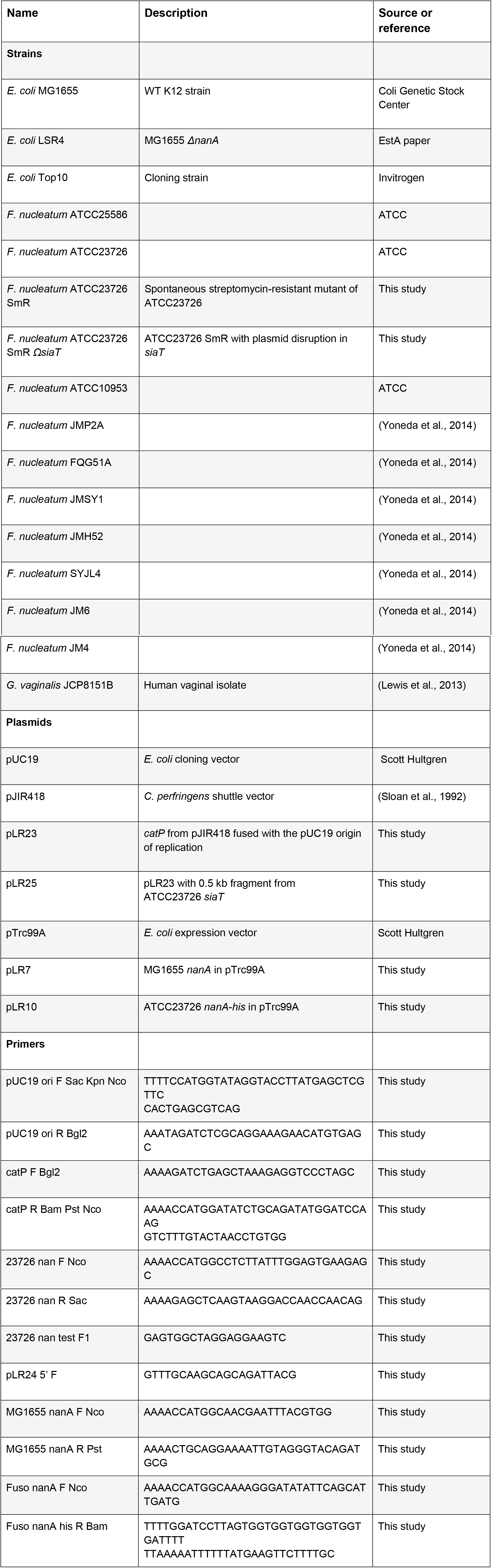
Key bacterial strains, plasmids and primers.

**Table S2.**
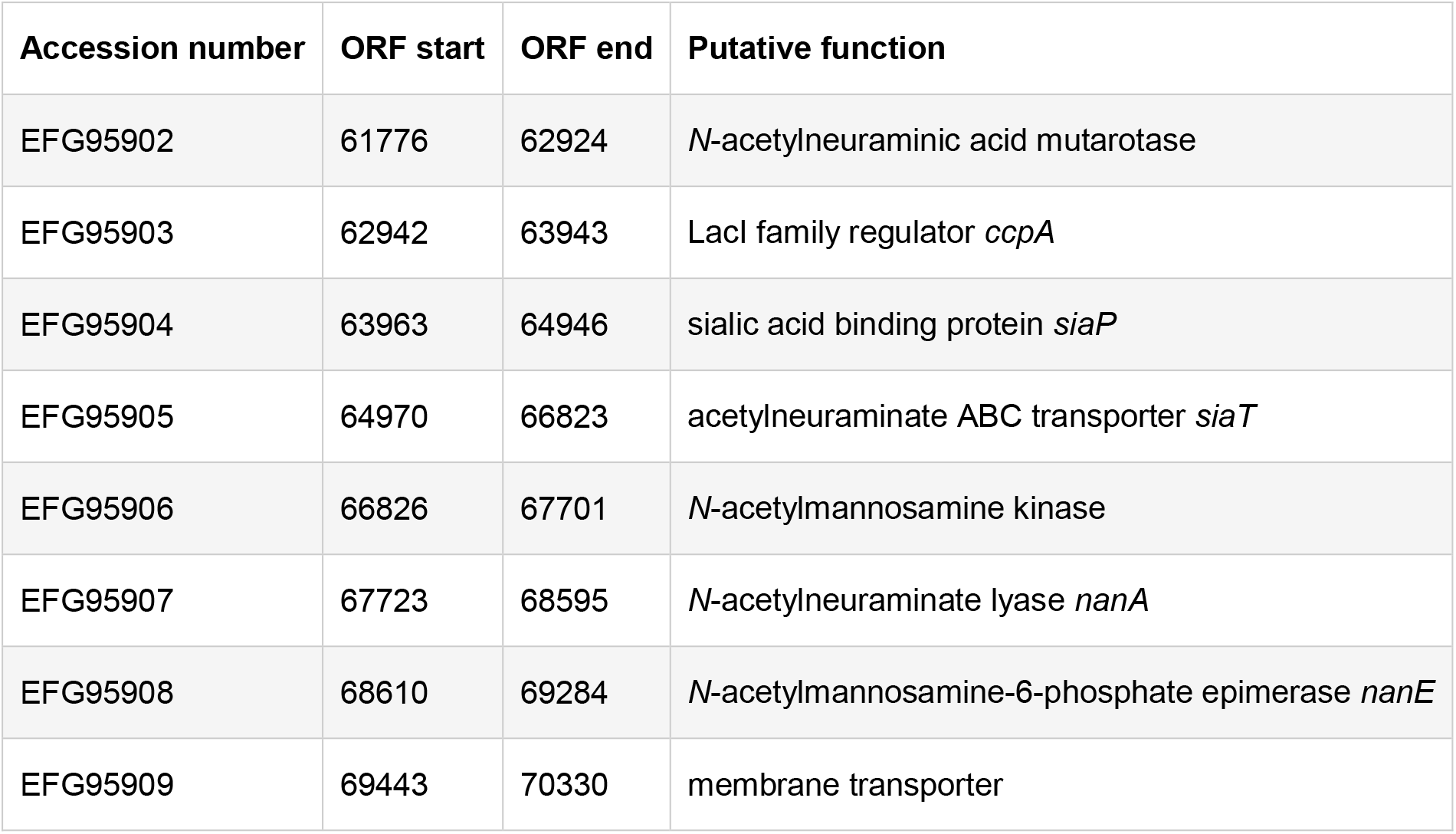
Genetic organization of the putative sialic acid catabolic gene cluster in *F. nucleatum* strain ATCC 23726, related to Figure 2.

| Accession number | ORF start | ORF end | Putative function |
| --- | --- | --- | --- |
| EFG95902 | 61776 | 62924 | *N*-acetylneuraminic acid mutarotase |
| EFG95903 | 62942 | 63943 | LacI family regulator *ccpA* |
| EFG95904 | 63963 | 64946 | sialic acid binding protein *siaP* |
| EFG95905 | 64970 | 66823 | acetylneuraminate ABC transporter *siaT* |
| EFG95906 | 66826 | 67701 | *N*-acetylmannosamine kinase |
| EFG95907 | 67723 | 68595 | *N*-acetylneuraminate lyase *nanA* |
| EFG95908 | 68610 | 69284 | *N*-acetylmannosamine-6-phosphate epimerase *nanE* |
| EFG95909 | 69443 | 70330 | membrane transporter |

**Figure S1.**
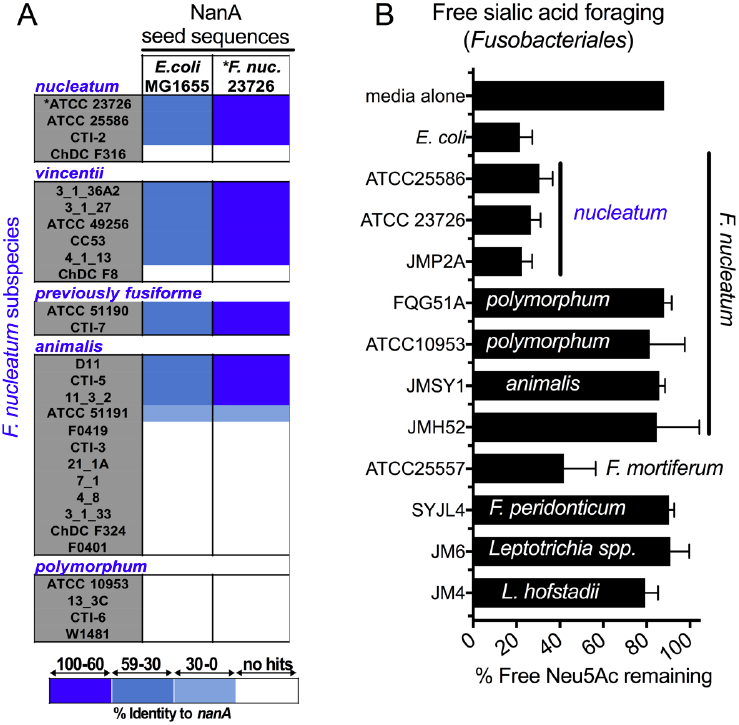
A sub group of *F. nucleatum* strains encode putative sialic acid lyase (NanA) homologs and consume free sialic acid, related to Figure 1. (A) Heat map showing percent identity of NanA homologs in sequenced *F. nucleatum* strains to the sialic acid lyase of *E. coli* MG1655. Homologs showed high similarity to amino acid sequence of lyase from *F. nucleatum 23726*. (B) Amount of sialic acid remaining in the growth medium 24-48 hours post inoculation of different *F. nucleatum* strains (as indicated), relative to un-inoculated media control. All bacteria were grown in medium supplemented with free sialic acid (Neu5Ac, 100 μM). Error bars shown are standard deviation of the mean, from three or more experiments.

**Figure S2.**
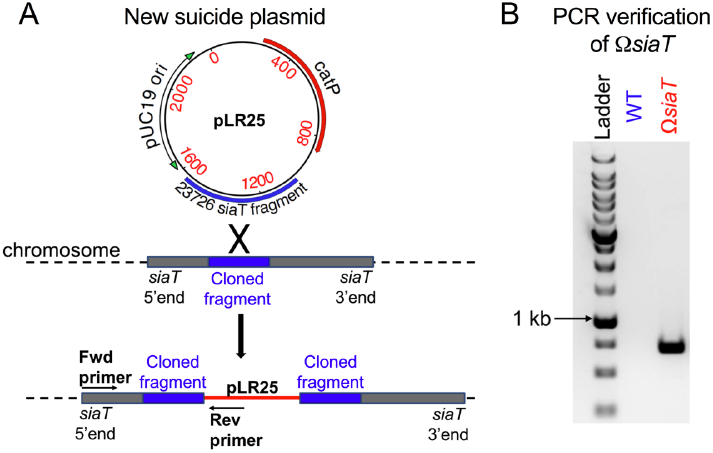
Insertion mutagenesis to disrupt sialic acid catabolic pathway in *F. nucleatum*, related to Figure 1. (A) Schematic showing integration of plasmid containing *siaT* insert (pLR25) into the *F. nucleatum* chromosome. Positions of forward (Fwd) and reverse (Rev) primers used for confirming integration of plasmid are also indicated. (B) Agarose gel image with the expected PCR product confirming integration of plasmid into the *siaT* locus.

**Figure S3.**
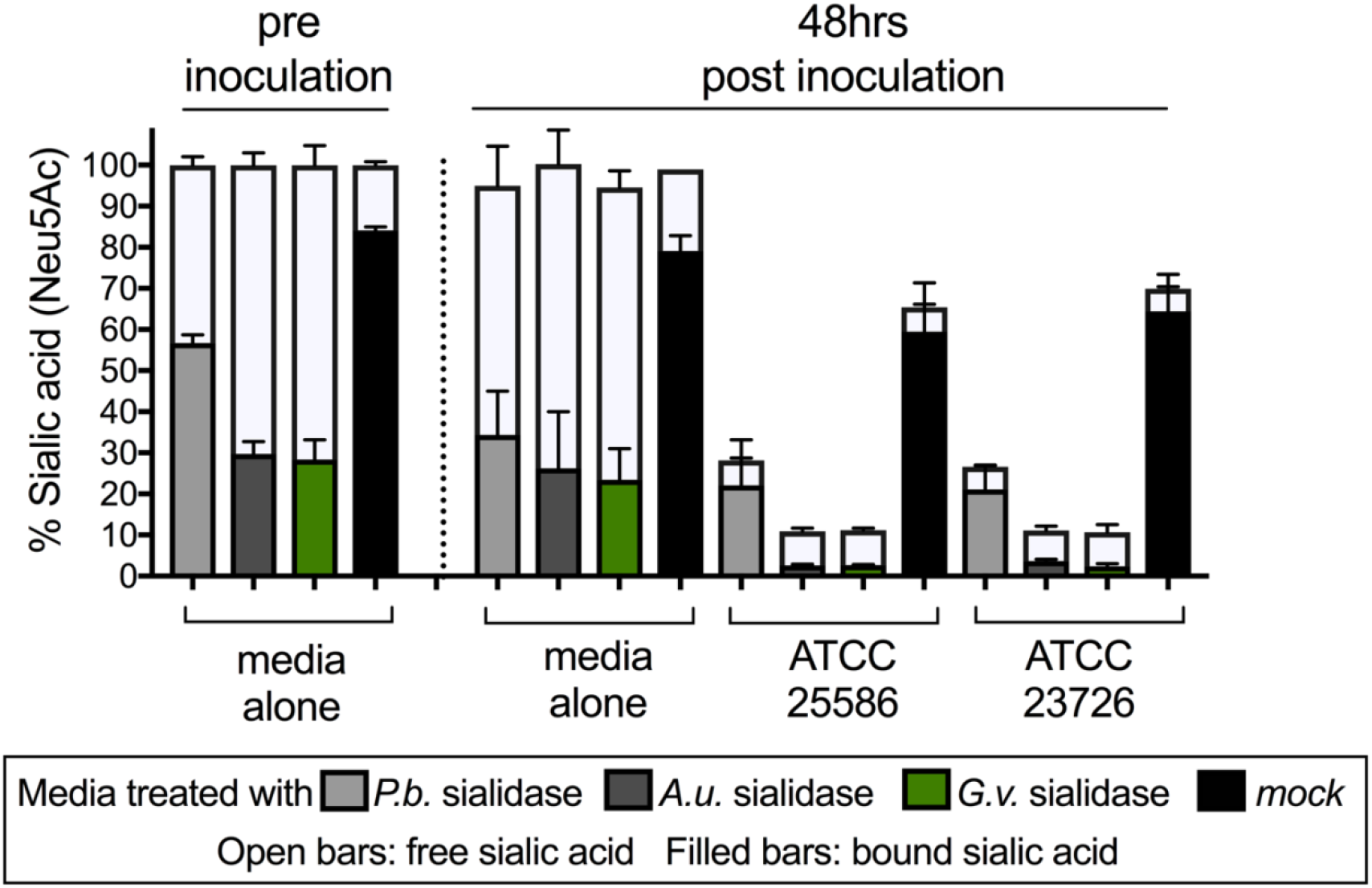
*F. nucleatum* ATCC23726 uptakes free sialic acid released by *P. bivia* sialidases, related to Figure 1. Relative amount of bound (bound = total-free) and free sialic acid remaining in the medium at 0 and 48 hours post inoculation. *F. nucleatum* was grown anaerobically in media that was either untreated or exposed to purified sialidase from *Arthrobacter ureafaciens* (*A.u.*) or *P. bivia (P.b.)* ATCC 29303. Bound sialic acids are inaccessible to *F. nucleatum* except in the presence of exogenous sialidase. Error bars represent standard deviation of the mean value.

**Figure S4.**
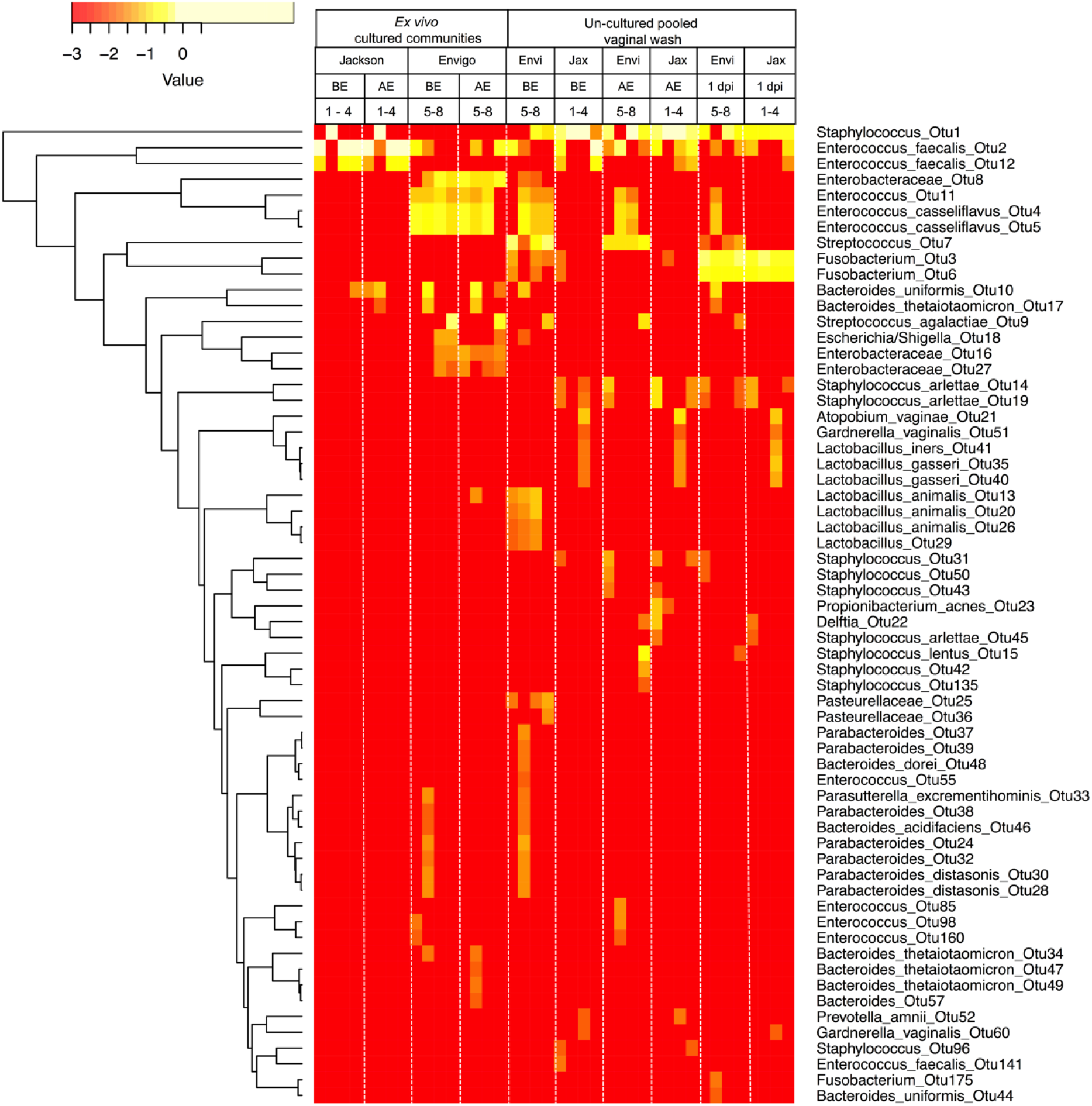
Vaginal microbiomes of C57BL/6 mice from Jackson and Envigo facilities, related to Figure 3. Heat map showing the log10 abundance of different OTUs in the microbiota pools of Jackson (Jax, 1-4) mice and Envigo (Envi, 5-8) collected before estrogenization (BE), after estrogenization (AE), and at 1 day post inoculation (dpi) with *F. nucleatum*. Microbiome analysis was done on uncultured and cultured vaginal washes pooled from 5 mice in the same cage. Each column represents one microbiota pool = 1 cage = 5 mice, total = 4 microbiota pools per vendor per condition. OTUs were assigned using the UNOISE algorithm. Data was clustered using hierarchical Euclidean clustering in the R project for statistical computing.

**Figure S5.**
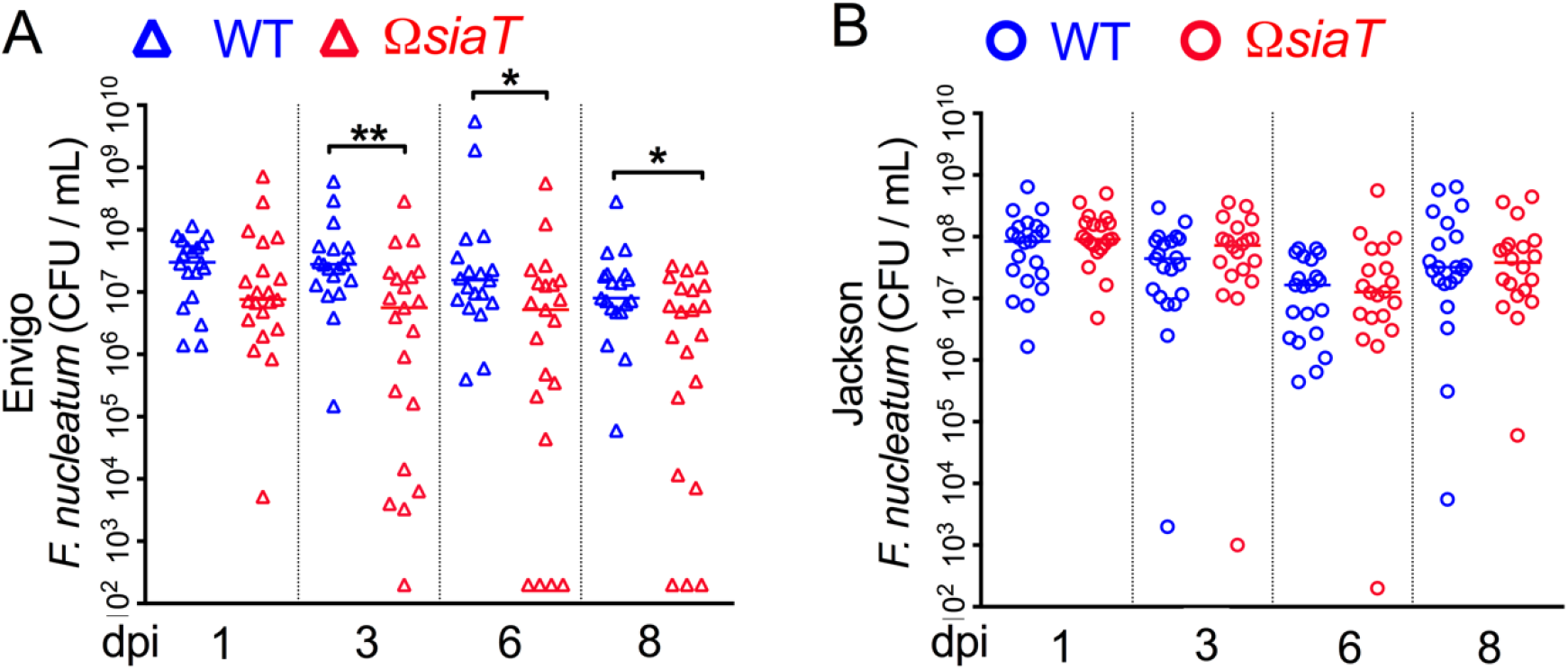
Vendor specific differences in *F. nucleatum* WT and *ΩsiaT* vaginal colonization in C57BL/6 mice, related to Figure 4. (A, B) *F. nucleatum* titers in vaginal wash collected from individual mice. (A) WT and *ΩsiaT* titers in mice from Envigo facility from 1 to 8 dpi. (B) WT and *ΩsiaT* titers in mice from Jackson facility from 1 to 8 dpi. Data is combined from 2 independent experiments, dpi = days post inoculation. Each experiment had 10 mice per group.

**Figure S6.**
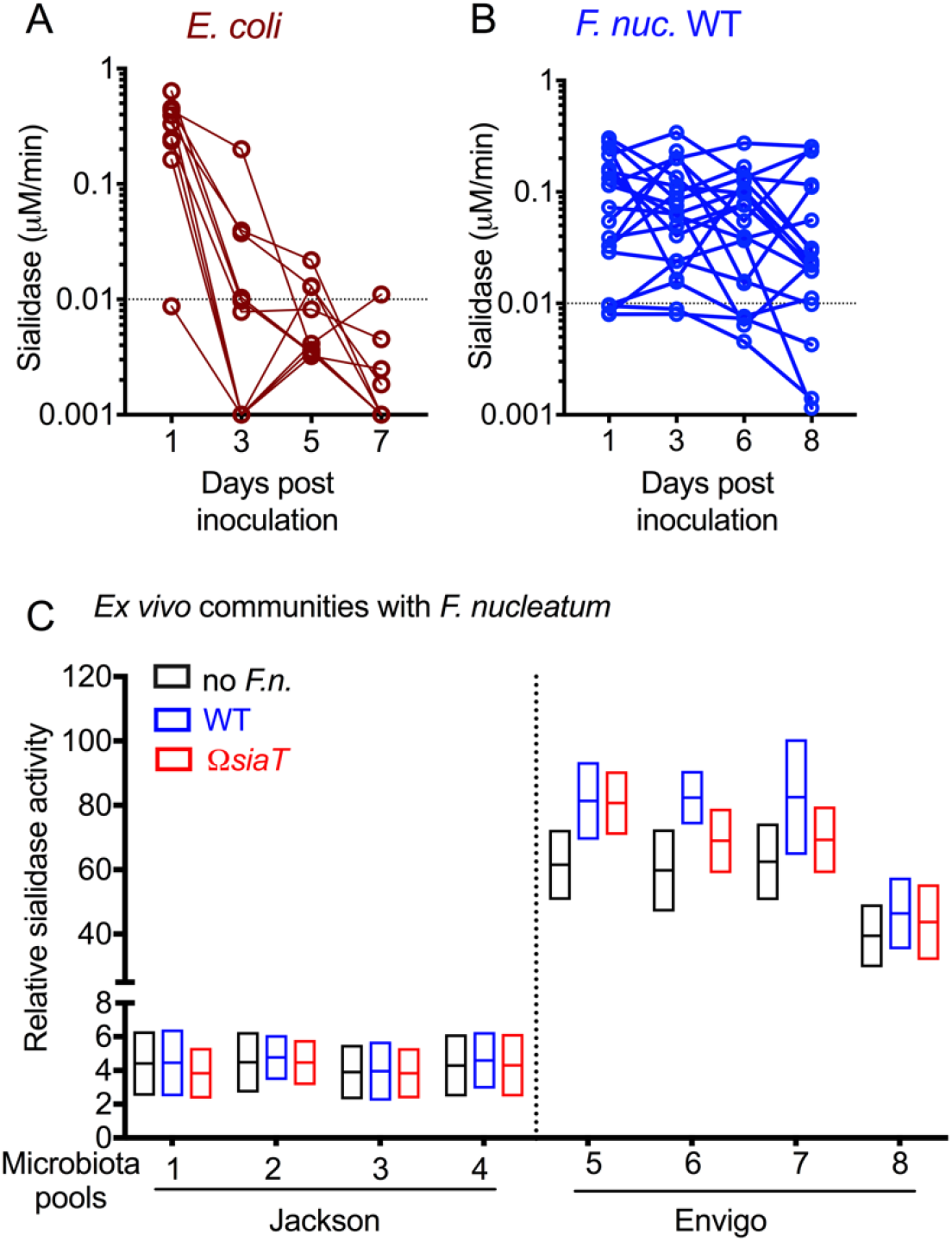
Vaginal sialidase activity in *E. coli* and *F. nucleatum* inoculated mice, and in vaginal microbiota pools of Jackson and Envigo mice, related to Figure 6. (A-B) Sialidase activity in vaginal washes from individual animals purchased from Envigo, estrogenized, and inoculated with *E. coli*, or *F. nucleatum* from 1 to 8 dpi. (C) Sialidase activity (after estrogenization) in vaginal wash pooled specimens (1 microbiota pool = 1 cage = 5 mice) from Jackson and Envigo mice, cultured with or without *F. nucleatum* (WT or *ΩsiaT*). No sialidase activity was observed in cultured vaginal microbial communities from Jackson mice, both with and without *F. nucleatum*. Data shown here is combined from 2 independent experiments.

## References

Allsworth, J.E., and Peipert, J.F. (2007). Prevalence of bacterial vaginosis: 2001-2004 National Health and Nutrition Examination Survey data. Obstet Gynecol 109, 114–120.

Altschul, S.F., Madden, T.L., Schaffer, A.A., Zhang, J., Zhang, Z., Miller, W., and Lipman, D.J. (1997). Gapped BLAST and PSI-BLAST: a new generation of protein database search programs. Nucleic Acids Res 25, 3389–3402.

Belzer, C., Chia, L.W., Aalvink, S., Chamlagain, B., Piironen, V., Knol, J., and de Vos, W.M. (2017). Microbial Metabolic Networks at the Mucus Layer Lead to Diet-Independent Butyrate and Vitamin B12 Production by Intestinal Symbionts. MBio 8.

Binder Gallimidi, A., Fischman, S., Revach, B., Bulvik, R., Maliutina, A., Rubinstein, A.M., Nussbaum, G., and Elkin, M. (2015). Periodontal pathogens *Porphyromonas gingivalis* and *Fusobacterium nucleatum* promote tumor progression in an oral-specific chemical carcinogenesis model. Oncotarget 6, 22613–22623.

Brennan, C.A., and Garrett, W.S. (2016). Gut Microbiota, Inflammation, and Colorectal Cancer. Annu Rev Microbiol 70, 395–411.

Briselden, A.M., Moncla, B.J., Stevens, C.E., and Hillier, S.L. (1992). Sialidases (neuraminidases) in bacterial vaginosis and bacterial vaginosis-associated microflora. J Clin Microbiol 30, 663–666.

Brotman, R.M., Klebanoff, M.A., Nansel, T.R., Yu, K.F., Andrews, W.W., Zhang, J., and Schwebke, J.R. (2010). Bacterial vaginosis assessed by gram stain and diminished colonization resistance to incident gonococcal, chlamydial, and trichomonal genital infection. The Journal of infectious diseases 202, 1907–1915.

Bullman, S., Pedamallu, C.S., Sicinska, E., Clancy, T.E., Zhang, X., Cai, D., Neuberg, D., Huang, K., Guevara, F., Nelson, T., et al. (2017). Analysis of *Fusobacterium* persistence and antibiotic response in colorectal cancer. Science 358, 1443–1448.

Caing-Carlsson, R., Goyal, P., Sharma, A., Ghosh, S., Setty, T.G., North, R.A., Friemann, R., and Ramaswamy, S. (2017). Crystal structure of *N*-acetylmannosamine kinase from *Fusobacterium nucleatum*. Acta Crystallogr F Struct Biol Commun 73, 356–362.

Cauci, S., and Culhane, J.F. (2011). High sialidase levels increase preterm birth risk among women who are bacterial vaginosis-positive in early gestation. American journal of obstetrics and gynecology 204, 142 e141–149.

Davis, G., and Gibbons, R.J. (1990). Accessible sialic acid content of oral epithelial cells from healthy and gingivitis subjects. J Periodontal Res 25, 250–253.

de Jonge, M.I., Keizer, S.A., El Moussaoui, H.M., van Dorsten, L., Azzawi, R., van Zuilekom, H.I., Peters, P.P., van Opzeeland, F.J., van Dijk, L., Nieuwland, R., et al. (2011). A novel guinea pig model of *Chlamydia trachomatis* genital tract infection. Vaccine 29, 5994–6001.

Denes, E., and Barraud, O. (2016). *Fusobacterium nucleatum* infections: clinical spectrum and bacteriological features of 78 cases. Infection 44, 475–481.

DiGiulio, D.B. (2012). Diversity of microbes in amniotic fluid. Semin Fetal Neonatal Med 17, 2–11.

Edgar, R.C. (2010). Search and clustering orders of magnitude faster than BLAST. Bioinformatics 26, 2460–2461.

Fukui, K., Fukui, Y., and Moriyama, T. (1971). Neuraminidase activity in some bacteria from the human mouth. Arch Oral Biol 16, 1361–1365.

Gangi Setty, T., Cho, C., Govindappa, S., Apicella, M.A., and Ramaswamy, S. (2014). Bacterial periplasmic sialic acid-binding proteins exhibit a conserved binding site. Acta Crystallogr D Biol Crystallogr 70, 1801–1811.

Gilbert, N.M., Lewis, W.G., and Lewis, A.L. (2013). Clinical features of bacterial vaginosis in a murine model of vaginal infection with *Gardnerella vaginalis*. PLoS One 8, e59539.

Goldenberg, R.L., Klebanoff, M.A., Nugent, R., Krohn, M.A., Hillier, S., and Andrews, W.W. (1996). Bacterial colonization of the vagina during pregnancy in four ethnic groups. Vaginal Infections and Prematurity Study Group. American journal of obstetrics and gynecology 174, 1618–1621.

Guerrero-Preston, R., White, J.R., Godoy-Vitorino, F., Rodriguez-Hilario, A., Navarro, K., Gonzalez, H., Michailidi, C., Jedlicka, A., Canapp, S., Bondy, J., et al. (2017). High-resolution microbiome profiling uncovers *Fusobacterium nucleatum*, *Lactobacillus gasseri/johnsonii*, and *Lactobacillus vaginalis* associated to oral and oropharyngeal cancer in saliva from HPV positive and HPV negative patients treated with surgery and chemo-radiation. Oncotarget 8, 110931–110948.

Haines-Menges, B.L., Whitaker, W.B., Lubin, J.B., and Boyd, E.F. (2015). Host Sialic Acids: A Delicacy for the Pathogen with Discerning Taste. Microbiol Spectr 3.

Han, Y.W., Shen, T., Chung, P., Buhimschi, I.A., and Buhimschi, C.S. (2009). Uncultivated bacteria as etiologic agents of intra-amniotic inflammation leading to preterm birth. J Clin Microbiol 47, 38–47.

Hill, G.B. (1998). Preterm birth: associations with genital and possibly oral microflora. Ann Periodontol 3, 222–232.

Hill, G.B., Eschenbach, D.A., and Holmes, K.K. (1984). Bacteriology of the vagina. Scand J Urol Nephrol Suppl 86, 23–39.

Hillier, S.L., Martius, J., Krohn, M., Kiviat, N., Holmes, K.K., and Eschenbach, D.A. (1988). A case-control study of chorioamnionic infection and histologic chorioamnionitis in prematurity. The New England journal of medicine 319, 972–978.

Hitti, J., Hillier, S.L., Agnew, K.J., Krohn, M.A., Reisner, D.P., and Eschenbach, D.A. (2001). Vaginal indicators of amniotic fluid infection in preterm labor. Obstet Gynecol 97, 211–219.

Holst, E., Goffeng, A.R., and Andersch, B. (1994). Bacterial vaginosis and vaginal microorganisms in idiopathic premature labor and association with pregnancy outcome. J Clin Microbiol 32, 176–186.

Huang, Y.L., Chassard, C., Hausmann, M., von Itzstein, M., and Hennet, T. (2015). Sialic acid catabolism drives intestinal inflammation and microbial dysbiosis in mice. Nat Commun 6, 8141.

Jerse, A.E. (1999). Experimental gonococcal genital tract infection and opacity protein expression in estradiol-treated mice. Infect Immun 67, 5699–5708.

Kook, J.K., Park, S.N., Lim, Y.K., Choi, M.H., Cho, E., Kong, S.W., Shin, Y., Paek, J., and Chang, Y.H. (2013). *Fusobacterium nucleatum* subsp. *fusiforme* Gharbia and Shah 1992 is a later synonym of *Fusobacterium nucleatum* subsp. *vincentii* Dzink et al. 1990. Curr Microbiol 66, 414–417.

Kurniyati, K., Zhang, W., Zhang, K., and Li, C. (2013). A surface-exposed neuraminidase affects complement resistance and virulence of the oral spirochaete Treponema denticola. Mol Microbiol 89, 842–856.

Lewis, A.L., Cao, H., Patel, S.K., Diaz, S., Ryan, W., Carlin, A.F., Thon, V., Lewis, W.G., Varki, A., Chen, X., et al. (2007). NeuA sialic acid *O*-acetylesterase activity modulates *O*-acetylation of capsular polysaccharide in group B Streptococcus. J Biol Chem 282, 27562–27571.

Lewis, A.L., Deitzler, G.E., Ruiz, M.J., Weimer, C., Park, S., Robinson, L.S., Hallsworth-Pepin, K., Wollam, A., Mitreva, M., and Lewis, W.G. (2016). Genome Sequences of 11 Human Vaginal Actinobacteria Strains. Genome Announc 4.

Lewis, A.L., Desa, N., Hansen, E.E., Knirel, Y.A., Gordon, J.I., Gagneux, P., Nizet, V., and Varki, A. (2009). Innovations in host and microbial sialic acid biosynthesis revealed by phylogenomic prediction of nonulosonic acid structure. Proc Natl Acad Sci U S A 106, 13552–13557.

Lewis, A.L., Hensler, M.E., Varki, A., and Nizet, V. (2006). The group B streptococcal sialic acid *O*-acetyltransferase is encoded by neuD, a conserved component of bacterial sialic acid biosynthetic gene clusters. J Biol Chem 281, 11186–11192.

Lewis, A.L., and Lewis, W.G. (2012). Host sialoglycans and bacterial sialidases: a mucosal perspective. Cellular microbiology.

Lewis, A.L., Nizet, V., and Varki, A. (2004). Discovery and characterization of sialic acid *O*-acetylation in group B Streptococcus. Proc Natl Acad Sci U S A 101, 11123–11128.

Lewis, W.G., Robinson, L.S., Gilbert, N.M., Perry, J.C., and Lewis, A.L. (2013). Degradation, foraging, and depletion of mucus sialoglycans by the vagina-adapted Actinobacterium *Gardnerella vaginalis*. J Biol Chem 288, 12067–12079.

Lewis, W.G., Robinson, L.S., Perry, J., Bick, J.L., Peipert, J.F., Allsworth, J.E., and Lewis, A.L. (2012). Hydrolysis of secreted sialoglycoprotein immunoglobulin A (IgA) in ex vivo and biochemical models of bacterial vaginosis. J Biol Chem 287, 2079–2089.

Li, C., Kurniyati, Hu, B., Bian, J., Sun, J., Zhang, W., Liu, J., Pan, Y., and Li, C. (2012). Abrogation of neuraminidase reduces biofilm formation, capsule biosynthesis, and virulence of *Porphyromonas gingivalis*. Infect Immun 80, 3–13.

Li, N., Ren, A., Wang, X., Fan, X., Zhao, Y., Gao, G.F., Cleary, P., and Wang, B. (2015). Influenza viral neuraminidase primes bacterial coinfection through TGF-beta-mediated expression of host cell receptors. Proc Natl Acad Sci U S A 112, 238–243.

Lim, B., Zimmermann, M., Barry, N.A., and Goodman, A.L. (2017). Engineered Regulatory Systems Modulate Gene Expression of Human Commensals in the Gut. Cell 169, 547–558 e515.

Mally, M., Shin, H., Paroz, C., Landmann, R., and Cornelis, G.R. (2008). *Capnocytophaga canimorsus*: a human pathogen feeding at the surface of epithelial cells and phagocytes. PLoS Pathog 4, e1000164.

Manjunath, L., Guntupalli, S.R., Currie, M.J., North, R.A., Dobson, R.C.J., Nayak, V., and Subramanian, R. (2018). Crystal structures and kinetic analyses of *N*-acetylmannosamine-6-phosphate 2-epimerases from *Fusobacterium nucleatum* and *Vibrio cholerae*. Acta Crystallogr F Struct Biol Commun 74, 431–440.

McGregor, J.A., French, J.I., Jones, W., Milligan, K., McKinney, P.J., Patterson, E., and Parker, R. (1994). Bacterial vaginosis is associated with prematurity and vaginal fluid mucinase and sialidase: results of a controlled trial of topical clindamycin cream. American journal of obstetrics and gynecology 170, 1048–1059; discussion 1059-1060.

Mobley, H.L., Green, D.M., Trifillis, A.L., Johnson, D.E., Chippendale, G.R., Lockatell, C.V., Jones, B.D., and Warren, J.W. (1990). Pyelonephritogenic *Escherichia coli* and killing of cultured human renal proximal tubular epithelial cells: role of hemolysin in some strains. Infect Immun 58, 1281–1289.

Moncla, B.J., Braham, P., and Hillier, S.L. (1990). Sialidase (neuraminidase) activity among gram-negative anaerobic and capnophilic bacteria. J Clin Microbiol 28, 422–425.

Moncla, B.J., Chappell, C.A., Mahal, L.K., Debo, B.M., Meyn, L.A., and Hillier, S.L. (2015). Impact of bacterial vaginosis, as assessed by nugent criteria and hormonal status on glycosidases and lectin binding in cervicovaginal lavage samples. PLoS One 10, e0127091.

Mulvey, M.A., Schilling, J.D., and Hultgren, S.J. (2001). Establishment of a persistent *Escherichia coli* reservoir during the acute phase of a bladder infection. Infect Immun 69, 4572–4579.

Myziuk, L., Romanowski, B., and Johnson, S.C. (2003). BVBlue test for diagnosis of bacterial vaginosis. J Clin Microbiol 41, 1925–1928.

Ng, K.M., Ferreyra, J.A., Higginbottom, S.K., Lynch, J.B., Kashyap, P.C., Gopinath, S., Naidu, N., Choudhury, B., Weimer, B.C., Monack, D.M., et al. (2013). Microbiota-liberated host sugars facilitate post-antibiotic expansion of enteric pathogens. Nature 502, 96–99.

Noecker, C., Eng, A., Srinivasan, S., Theriot, C.M., Young, V.B., Jansson, J.K., Fredricks, D.N., and Borenstein, E. (2016). Metabolic Model-Based Integration of Microbiome Taxonomic and Metabolomic Profiles Elucidates Mechanistic Links between Ecological and Metabolic Variation. mSystems 1.

Nonaka, H., Ishikawa, Y., Otsuka, M., Toda, K., Sato, M., and Nakamura, R. (1983). Purification and some properties of neuraminidase isolated from the culture medium of oral bacterium *Streptococcus mitis* ATCC 9811. J Dent Res 62, 792–797.

Patras, K.A., and Doran, K.S. (2016). A Murine Model of Group B Streptococcus Vaginal Colonization. J Vis Exp.

Peipert, J.F., Lapane, K.L., Allsworth, J.E., Redding, C.A., Blume, J.D., and Stein, M.D. (2008). Bacterial vaginosis, race, and sexually transmitted infections: does race modify the association? Sex Transm Dis 35, 363–367.

Rakoff-Nahoum, S., Foster, K.R., and Comstock, L.E. (2016). The evolution of cooperation within the gut microbiota. Nature 533, 255–259.

Ravel, J., Gajer, P., Abdo, Z., Schneider, G.M., Koenig, S.S., McCulle, S.L., Karlebach, S., Gorle, R., Russell, J., Tacket, C.O., et al. (2011). Vaginal microbiome of reproductive-age women. Proc Natl Acad Sci U S A 108 Suppl 1, 4680–4687.

Rey, F.E., Gonzalez, M.D., Cheng, J., Wu, M., Ahern, P.P., and Gordon, J.I. (2013). Metabolic niche of a prominent sulfate-reducing human gut bacterium. Proc Natl Acad Sci U S A 110, 13582–13587.

Rezeberga, D., Lazdane, G., Kroica, J., Sokolova, L., and Donders, G.G. (2008). Placental histological inflammation and reproductive tract infections in a low risk pregnant population in Latvia. Acta Obstet Gynecol Scand 87, 360–365.

Robinson, L.S., Lewis, W.G., and Lewis, A.L. (2017). The sialate *O*-acetylesterase EstA from gut Bacteroidetes species enables sialidase-mediated cross-species foraging of 9-*O*-acetylated sialoglycans. J Biol Chem 292, 11861–11872.

Sakamoto, H., Kato, H., Sato, T., and Sasaki, J. (1998). Semiquantitative bacteriology of closed odontogenic abscesses. Bull Tokyo Dent Coll 39, 103–107.

Severi, E., Hood, D.W., and Thomas, G.H. (2007). Sialic acid utilization by bacterial pathogens. Microbiology 153, 2817–2822.

Siegel, S.J., Roche, A.M., and Weiser, J.N. (2014). Influenza promotes pneumococcal growth during coinfection by providing host sialylated substrates as a nutrient source. Cell Host Microbe 16, 55–67.

Silver, H.M., Sperling, R.S., St Clair, P.J., and Gibbs, R.S. (1989). Evidence relating bacterial vaginosis to intraamniotic infection. American journal of obstetrics and gynecology 161, 808–812.

Siqueira, J.F., Jr., Rocas, I.N., Souto, R., Uzeda, M., and Colombo, A.P. (2001). Microbiological evaluation of acute periradicular abscesses by DNA-DNA hybridization. Oral Surg Oral Med Oral Pathol Oral Radiol Endod 92, 451–457.

Sloan, J., Warner, T.A., Scott, P.T., Bannam, T.L., Berryman, D.I., and Rood, J.I. (1992). Construction of a sequenced *Clostridium perfringens*-*Escherichia coli* shuttle plasmid. Plasmid 27, 207–219.

Song, Y.G., Shim, S.G., Kim, K.M., Lee, D.H., Kim, D.S., Choi, S.H., Song, J.Y., Kang, H.L., Baik, S.C., Lee, W.K., et al. (2014). Profiling of the bacteria responsible for pyogenic liver abscess by 16S rRNA gene pyrosequencing. J Microbiol 52, 504–509.

Srinivasan, S., Morgan, M.T., Fiedler, T.L., Djukovic, D., Hoffman, N.G., Raftery, D., Marrazzo, J.M., and Fredricks, D.N. (2015). Metabolic signatures of bacterial vaginosis. MBio 6.

Tailford, L.E., Owen, C.D., Walshaw, J., Crost, E.H., Hardy-Goddard, J., Le Gall, G., de Vos, W.M., Taylor, G.L., and Juge, N. (2015). Discovery of intramolecular trans-sialidases in human gut microbiota suggests novel mechanisms of mucosal adaptation. Nat Commun 6, 7624.

Urushiyama, D., Suda, W., Ohnishi, E., Araki, R., Kiyoshima, C., Kurakazu, M., Sanui, A., Yotsumoto, F., Murata, M., Nabeshima, K., et al. (2017). Microbiome profile of the amniotic fluid as a predictive biomarker of perinatal outcome. Sci Rep 7, 12171.

Vimr, E.R., and Troy, F.A. (1985). Identification of an inducible catabolic system for sialic acids (nan) in *Escherichia coli*. J Bacteriol 164, 845–853.

Wang, L., Koppolu, S., Chappell, C., Moncla, B.J., Hillier, S.L., and Mahal, L.K. (2015). Studying the effects of reproductive hormones and bacterial vaginosis on the glycome of lavage samples from the cervicovaginal cavity. PLoS One 10, e0127021.

Wiesenfeld, H.C., Hillier, S.L., Krohn, M.A., Landers, D.V., and Sweet, R.L. (2003). Bacterial vaginosis is a strong predictor of *Neisseria gonorrhoeae* and *Chlamydia trachomatis* infection. Clin Infect Dis 36, 663–668.

Wong, A., Grau, M.A., Singh, A.K., Woodiga, S.A., and King, S.J. (2018). Role of Neuraminidase-Producing Bacteria in Exposing Cryptic Carbohydrate Receptors for *Streptococcus gordonii* Adherence. Infect Immun 86.

Yoneda, S., Loeser, B., Feng, J., Dmytryk, J., Qi, F., and Merritt, J. (2014). Ubiquitous sialometabolism present among oral fusobacteria. PLoS One 9, e99263.

Zhang, S., Cai, S., and Ma, Y. (2018). Association between *Fusobacterium nucleatum* and colorectal cancer: Progress and future directions. J Cancer 9, 1652–1659.

